# BIN1 genetic risk factor for Alzheimer is sufficient to induce early structural tract alterations in entorhinal-hippocampal area and memory-related hippocampal multi-scale impairments

**DOI:** 10.1101/437228

**Authors:** R Daudin, D Marechal, R Golgolab, Q Wang, Y Abe, T. Tsurugizawa, N Bourg, M Sartori, Y Loe-Mie, J Lipecka, C Guerrera, A McKenzie, B Potier, P Dutar, J Viard, A.M Lepagnol-Bestel, A Winkeler, I. Uszynski, V Hindié, MC Birling, L Lindner, C Chevalier, G Pavlovic, M Reis, H Kranz, G Dupuis, S Lévêque-Fort, J Diaz, E Davenas, D Dembele, H. Atas-Ozcan, J Laporte, C Thibault-Carpentier, B Malissen, J.C Rain, C. Poupon, D Le Bihan, B Zhang, Y Herault, L. Ciobanu, M Simonneau

## Abstract

Genetic factors are known to contribute to Late Onset Alzheimer’s disease (LOAD) but their contribution to pathophysiology, specially to prodomic phases accessible to therapeutic approaches are far to be understood.

To translate genetic risk of Alzheimer’s disease (AD) into mechanistic insight, we generated transgenic mouse lines that express a ∼195 kbp human BAC that includes only *BIN1*, a gene associated to LOAD. This model gives a modest BIN1 overexpression, dependent of the number of BAC copies. At 6 months of age, we detected impaired entorhinal cortex (EC)-hippocampal pathways with specific impairments in EC-dentate gyrus synaptic long-term potentiation, dendritic spines of granular cells and recognition episodic memory. Structural changes were quantified using MRI. Their whole-brain functional impact were analyzed using resting state fMRI with a hypoconnectivity centered on entorhinal cortex.

These early phenotype defects independent of any changes in A-beta can be instrumental in the search for new AD drug targets.

---

Late-onset Alzheimer Disease (LOAD) is the most common form of dementia and one of the most challenging diseases of modern society (Querfurth and LaFerla, 2010). Understanding the preclinical stages of AD that begins in the brain at least 2–3 decades before evidence of episodic memory defects in patients is pivotal for the design of successful approaches to delay or reverse the transition from normal brain function to cognitive impairments.

Our working hypothesis is that LOAD genetic risk factors can be sufficient to generate early phenotypical changes before any changes in either Abeta or Tau. We focused our interest on *BIN1* that is the second important risk factor for AD, following the APOE gene (Lambert et al., 2013) and generated an *hBIN1* mouse model based on a slight human *BIN1* gene overexpression that we found in post-mortem brain samples from LOAD patients.

The *B*ridging *IN*tegrator-1 (BIN1; also referred to as Amphiphysin 2) is part of the Bin/Amphiphysin/Rvs (BAR) family of adaptor proteins that regulates membrane dynamics in a variety of cellular functions. The BIN1 paralog Amphiphysin 1 has a strict neuronal expression and presynaptic localization. Based on its sequence similarity, it has been proposed that BIN1 might function in synaptic vesicle endocytosis but *Bin1*^−/−^ mice failed to reveal a deficiency in basal synaptic vesicle uptake raising the possibility that BIN1 and Amphiphysin 1 may have divergent functions in the brain (Karch et al., 2014; Prokic et al., 2014).

The LOAD-associated *BIN1* SNP location 28 kb upstream of the *BIN1* coding region (Lambert et al., 2013) suggests that altered BIN1 expression might constitute the genetic mechanism by which *BIN1* increases LOAD risk (Chapuis et al., 2013). We first analyzed BIN1 expression changes found in postmortem brain tissues of LOAD patients. We analyzed 8 datasets of BIN1 transcripts levels (Zhang et al., 2013) in the human brain of either LOAD patients or controls and observed increased BIN1 transcripts levels in brains of AD cases when compared with controls (Fig. 1A).

**Figure 1.**
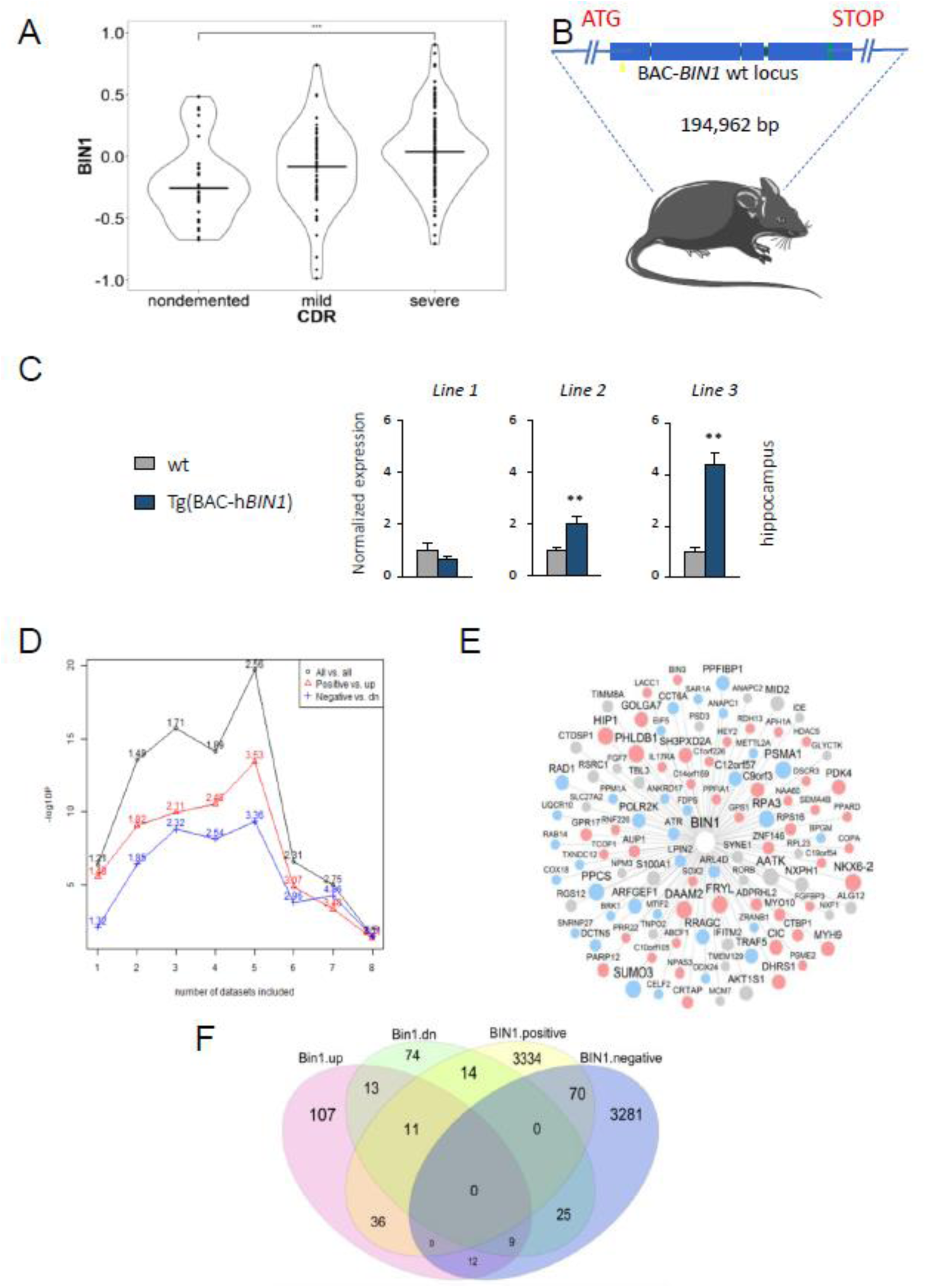
Seven-week-old *hBIN1* mouse hippocampus subregions share co-regulated genes with postmortem human brain subregions of patients with Alzheimer disease. **a.** BIN1 is significantly upregulated in the Brodmann area 36 (BM36) that is part of the perirhinal cortex in Alzheimer’s Disease patients with severe dementia compared with non-demented controls (1.2 fold change, corrected p=6.5E-4). **b.** Schematic representation of hBIN1 mouse line **c. c.** Normalized expression of murine of Bin1 in quadriceps and brain of both wt and different Tg extracts (Line 1: U154.20.8, Line2: U154.45.12; Line3: U154.16.16,). All percentage ratios were based on Bin1 RNA (left) and protein (right) level in wt samples. * p<0.05; ** p<0.01; *** p < 0.001. **d.** Enrichment of the DEG signature from *Bin1* transgenic mouse brain with the consensus *BIN1*-correlated gene signatures CCGS(*x*), where, x=2, 3, …, 8. Number on the top of each dot represents the fold enrichment. Black line indicates the overall enrichment regardless of correlation direction. Red line indicates the intersection between the gene signatures that are positively correlated with *BIN1* in human brain and the up-regulated genes in *Bin1* transgenic mice while the blue line indicates the enrichment between those negatively correlated with *BIN1* and the down-regulated genes in the same setting. The height datasets used to generate Bin1-centered network are from Mount Sinai Brain Bank (MSBB) which contains data from 4 brain regions (BM10,22,36,44), Harvard brain bank (HBTRC) which contains data from 3 brain regions (PFC, VC and CR) and ROSMAP data which only has 1 region. **e.** A *BIN1*-centered network confirmed in both the human AD brain studies (using CCGS(5)) and the *hBIN1* transgenic mice. Node size is proportional to the number of datasets in which a gene is significantly correlated with *BIN1*. Red nodes are the genes that are positively correlated with *BIN1* in human AD studies and are up-regulated in *Bin1* transgenic mouse model. Blue nodes are the genes negatively correlated with *BIN1* in the human AD studies and are down-regulated in Bin-1 transgenic mouse model. Grey nodes are the genes significantly correlated with *BIN1* but in different directions across multiple datasets. **f.** Venn diagram of the DEGs from *Bin1*-transgenic mice and *BIN1*-correlated signatures from

Taking these expression data in account, we generated eight distinct transgenic mouse lines based on a slight *BIN1* overexpression, using a 194,962 bp human BAC that includes exclusively the *BIN1* gene (Fig. 1B) (Supplementary Figures 1-2).We focused our analysis on three lines that incorporated different number of BAC copies: Line 1: U154.20.8 (3 copies), Line2: U154.45.12 (4-7 copies) and Line3: U154.16.16 (5-10 copies) (Supplementary Figure 3). We found that these lines express hBIN1 in mouse hippocampus in relation with the number of incorporated BACs (Fig.1C).

We next compared differentially expressed genes (DEG) in the hippocampus of this *hBIN1* mouse model with DEG repertoires found in post-mortem samples of LOAD and controls, expecting to find a repertoire of genes deregulated by BIN1 overexpression and common to both mouse and human. We used laser-assisted microdissection of 7 week-old mouse hippocampus subregions (Dorsal and ventral CA1/2, CA3 and Dentate Gyrus) from sagittal brain slices (n=5 WT; n=5 *hBIN1*). By the cutoff of fold change >1.2 (or <0.833) and BH-corrected p value <0.2, we identified 690 up-regulated and 1016 down-regulated genes in the *hBIN1* mouse, among which many were microRNAs (Supplementary Table 1). The down-regulated genes were most significantly enriched with olfactory receptor activity with corrected FET p=2.5E-16 (4.7 fold enrichment) while the up-regulated genes were not significantly enriched with any known pathways. To address to what extent Bin1 transgenic mouse model can represent AD pathogenesis and progression in human, we identified BIN1-correlated gene signatures for constructing a BIN1-centered correlation network from 8 datasets in 3 independent studies of human AD postmortem brains. We then built up consensus BIN1-correlated gene signatures CCG(x) in which each gene is significantly correlated with BIN1 in at least x datasets (2≤x≤8). The significance of the overlap between CCGS (x) and the differentially expressed gene (DEG) signatures from *hBIN1* transgenic mouse model was examined by FET. CCGS (*5*) is most significantly enriched for the DEG signature (Fig. 1D). In CCGS(*5*), the genes negatively correlated with *BIN1* were involved in synaptic transmission (corrected FET p=8.4E-23 and 1.8 FE) and mitochondria-related functions such as oxidative phosphorylation (corrected FET p=1.0E-18 and 2.9 FE) and respiratory electron transport chain (corrected FET p=7.0E-22 and 3.1 FE), suggesting a potential synaptic dysfunction and disruption of energy homeostasis while the genes positively correlated with BIN1 were associated with I-kappaB kinase/NF-kappaB signaling (corrected FET p=1.0E-04 and 1.5 FE), gliogenesis (corrected FET p=2.0E-04 and 1.6 FE) and intrinsic apoptotic signaling pathway to DNA damages (corrected FET p=1.6E-04 and 1.9 FE), indicating the increase inflammatory responses and cell death with Bin1 overexpression (Supplementary Tables 2, 3). We further examined the intersection of consensus *BIN1*-correlated gene signatures with the DEG signatures from *hBIN1* transgenic mouse brain (Fig. 1 E, F) (Supplementary Table 4). This network showed the core targets of *BIN1* as the expression of each gene was not only induced by *BIN1* in *hBIN1* transgenic mouse model but also correlated with the expression of *BIN1* in human AD brains. The analysis of the common repertoires between mouse *hBIN1* 7-week mouse hippocampus sub-regions hippocampus and LOAD brains highlighted the downregulation of chemical synaptic transmission gene repertoire and the upregulation of gliogenesis and inflammation-related repertoires (Supplementary Tables 2 and 3). Altogether, the identification of co-deregulated gene repertoires common to mouse *hBIN1* hippocampus and post-mortem brain samples from LOAD patients demonstrates the validity of the *hBIN1* model.

We next examined whether structural tract alterations can be evidenced in *hBIN1* mice as a possible consequence of these molecular deregulations. We used established diffusion tensor imaging (DTI) methods (Jones and Leemans, 2011) taking advantage of an ultra-high field (17.2Tesla) Magnetic Resonance Imaging (MRI) system. We studied the three distinct hBIN1 lines described in Fig. 1C. We used an analysis of mean diffusivity (MD) focusing on Regions of Interest (ROIs) within the hippocampal-entorhinal area (CA1, CA3, DG, Entl, Entm, Sub), and within the whole brain (Au, Hippo, MOp, SS, Spt, Tea, VISa, mPFC, Ent, RSP). Line 1 that do not express hBIN1 in mouse hippocampus did not display any structural changes (Supplementary Figure 4). In contrast, for lines 2 and 3, MD DTI scalars changes were evidenced at month 3 in medial entorhinal cortex and lateral septum (Fig. 2A-B). Interestingly, at month 6, we found a spreading of these structural defects in the hippocampal-entorhinal cortex complex with all the structures impacted and in primary sensory cortex (auditory and somatosensory cortexes) (Fig. 2C). Entorhinal Cortex is the first structure to display tau tangles in AD (Braak and Braak, 1995; Braak et al., 1993) with functional defects starting in Lateral Entorhinal Cortex (LEC) (Hyman et al., 1984; Khan et al., 2014b; Small, 2014b). Entorhinal–hippocampal neuronal circuits are crucial for spatial and contextual representation, navigation, and episodic memory (O’Keeffe and Nadel, 1978). We also identified structural tract alterations in the lateral septum region (Fig. 2A, B). The lateral septum (LS) is a major target of the hippocampal CA3 and CA1 pyramidal neurons, with a density of hippocampal afferents is ∼20-fold higher when compared to entorhinal regions or ∼180-fold higher when compared to prefrontal targets (Tingley and Buzsáki, 2018). This makes the LS the recipient of massive convergent excitation from the hippocampal-subicular-entorhinal population. This major anatomical connection convey hippocampal activity to its hypothalamic, mesencephalic, and brainstem projections (Tingley and Buzsáki, 2020).

**Figure 2.**
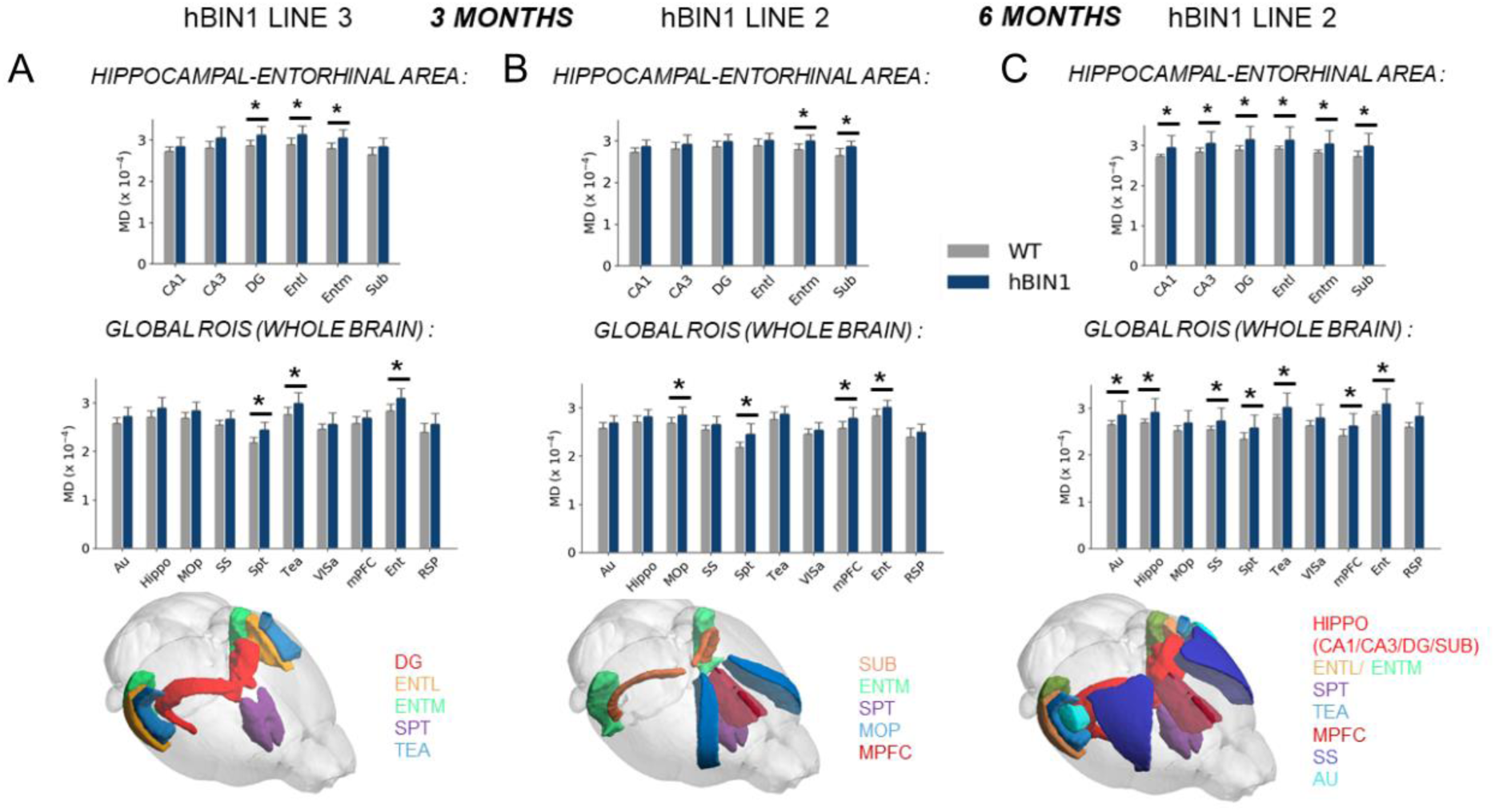
Whole-brain MRI-DTI analysis: MD increases in *hBIN1* transgenic mice. MD values for *hBIN1* and wild-type control (WT) mice, at 3 months (*hBIN1* lines 2 and 3) and 6 months (line 2). Two sets of ROIs are analyzed: within the hippocampal-entorhinal area (CA1, CA3, DG, Entl, Entm, Sub), and within the whole brain (Au, Hippo, MOp, SS, Spt, Tea, VISa, mPFC, Ent, RSP). We show a spreading of the abnormalities from 3 months (**A, B**) to 6 months (**C**) within the whole hippocampal-entorhinal area as well as other cortical areas. The adjusted p-values were obtained using Tukey HSD statistical tests with significance level alpha=0.05. Under the graphs are represented the significant ROIs on a 3D mouse brain. *p<0.05 Abbreviations: Au : primary auditory area ; Hippo : hippocampus ; MOp : primary motor area ; SS : somatosensory area ; Spt : lateral septal nucleus ; Tea : temporal association areas ; VISa : visual areas ; mPFC : medial prefontal cortex ; Ent : entorhinal cortex ; RSP : retrosplenial cortex ; DG : dentate gyrus ; Entl : entorhinal area, lateral part ; Entm : entorhinal area, medial part ; Sub : subiculum

Taking in account these EC-hippocampal sub-regions structural tract alterations, we analyzed possible related cellular phenotypes. We first focused on the analysis of dendritic spines in relevant neurons. We found an increase in filopodia and a decrease in mature spines of neurons from the dorsal DG, using diI biolistic injection of hippocampal slices at 3 months (Fig. 3A, B). This result demonstrates an impairment of the number of functional spines in dendrites of DG neurons in *hBIN1* mice, possibly linked to changes in entorhinal cortex-dentate gyrus pathway (Fig. 3C). From these results on dendritic spine impairment, we next analyzed if dendrites of DG neurons displayed structural changes. Using Sholl analysis, we quantified the number of neuronal branches to virtual build the dendritic tree and observed a statistically significant dendritic simplification in DG neurons (Supplementary Figure 5). We further examined synaptic transmission and plasticity in hippocampal slices at 3 months. When we stimulated the EC Perforant Path (PP) and recorded population responses in the dorsal dentate gyrus, we evidenced an impaired long-term potentiation in *hBIN1* mice (Fig. 3D). In contrast, analysis of synapses between CA3 and CA1 indicated no changes in long-term potentiation (LTP) in *hBIN1* mice (Fig. 3E). Together, these results indicate that *hBIN1* mice display an early contrasted synaptic plasticity phenotype limited to EC-DG synapses.

**Figure 3.**
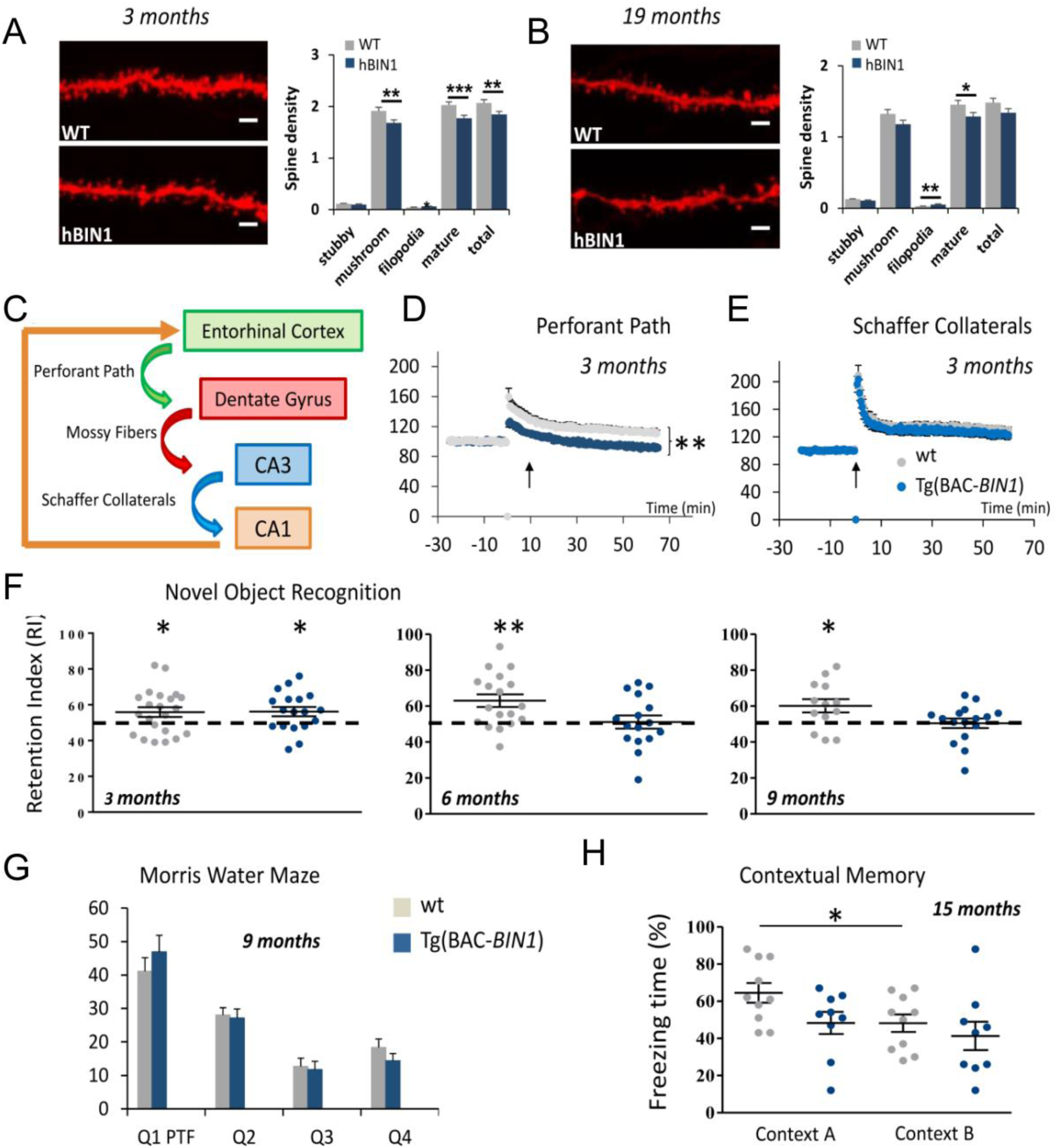
Early phenotype centered on impact on Entorhinal Cortex-Dentate Gyrus at month 3 and its episodic memory consequences, followed by a spreading of this phenotype at month 15. **(A)** Impact on spine density of dorsal dentate gyrus granular cells using DiI labelling of coronal slices in wt and hBIN1 mice at month 3. **(B)** Impact on spine density of dorsal dentate gyrus granular cells using DiI labelling of coronal slices in wt and hBIN1 mice at month 19. For each, n=3 wt; n=3 hBIN1 for > 8 dendrites per animal. Scale bar=2,5µm. * p < 0.05; ** p<0.005; *** p < 0.0005. **(C)** Schematic representation of entorhinal-hippocampal connections **(D)** Specific impact on synaptic plasticity. Theta-burst stimulation of the PP input to the DG induced potentiation of fEPSP in *control* (grey circles) but not in transgenic (blue circles). **(E)** In contrast, theta-burst stimulation of the CA3 input to the CA1 (Schaffer collaterals) induced normal potentiation of fEPSP in *control* (grey circles) and transgenic (blue circles). Error bars are the SEMs. **(F)** Novel Object Recognition memory is altered in transgenic human Bin1 mice from 6 months. The discrimination index (see Methods) is impaired at 6 and 9 months. (* p<0.05 and ** p<0.01). **(G)** Spatial reference episodic memory is not affected in transgenic BIN1 mice (tested at 9 months). Both wt and Tg mice spent more time in the reference (Q1 PFT) quadrant compared to hazard (25%) (*One sample t-test, ** p<0,01)*. Similar results have been obtained in 15 month-old *hBIN1* mice. **(H)** Pattern recognition of context episodic memory is altered in transgenic human Bin1 mice at 15 months (see M&M for paradigm description). Transgenic individuals displayed higher freezing time compared to controls (fear conditioning test) (*Statistics: Student t-test, * p<0,05, ** p<0,01)*.

We further studied if these structural and functional defects can induce *in vivo* hippocampus related-cognitive changes. We were able to identify such changes at month 6. We found that *hBIN1* mice showed a deficit in novel object recognition at months 6 and 9 (Fig. 3F) (Supplementary Figure 6). In contrast, spatial long-term memory using Morris Water Maze was found normal at months 9 (Fig. 3G) (Supplementary Figure 7). There is a converging consensus that the hippocampus supports episodic memory by combining a spatial/temporal “where” signal (from the MEC) with a “what” signal related to individual items experienced by the animal (from the LEC) (Connor and Knierim, 2017). Here, we evidenced a contrasted impairment in object recognition with no changes in spatial recognition. Altogether these results suggest that the *hBIN1* early phenotype is linked to a specific impact on LEC-DG pathway that is the earliest affected both in human aging (Reagh et al., 2018) and in AD (Braak and Braak, 1995; Braak et al., 1993; Khan et al., 2014a; Small, 2014a).

We analyzed if this early phenotype was worsening with time as expected for a preclinical model of neurodegeneration and found a progression of structural tract alterations from month 3 to month 6 using MRI-DTI (Fig.2). Alterations in MD at month 6 include primary sensory cortices that are known to be directly connected to LEC (Zingg et al., 2014).

We next investigated whole-brain functional connectivity of hBIN1 as a function of time in line 3 (Fig. 4A). Resting state functional magnetic resonance imaging (rs-fMRI) is used in brain mapping to evaluate regional interactions that occur in a resting state, when an explicit task is not being performed (Buckner et al., 2013). At month 6, we evidenced a mixed hypo-connectivity and hyper-connectivity for regions linked to episodic memory (CA1, LEC, RSP, mPFC, Lateral Septum, TEA) (Fig. 4B). We identified in particular a hypo-connectivity between CA1, LEC, mPFC and Lateral Septum This hypo-connectivity centered on LEC at month 6 is compatible with functional impairments reported in patients displaying minor cognitive impairment (MCI) (Sorg et al., 2007) and suggests that whole-brain functional activity of *hBIN1* mice mimics that found in MCI patients.

**Figure 4.**
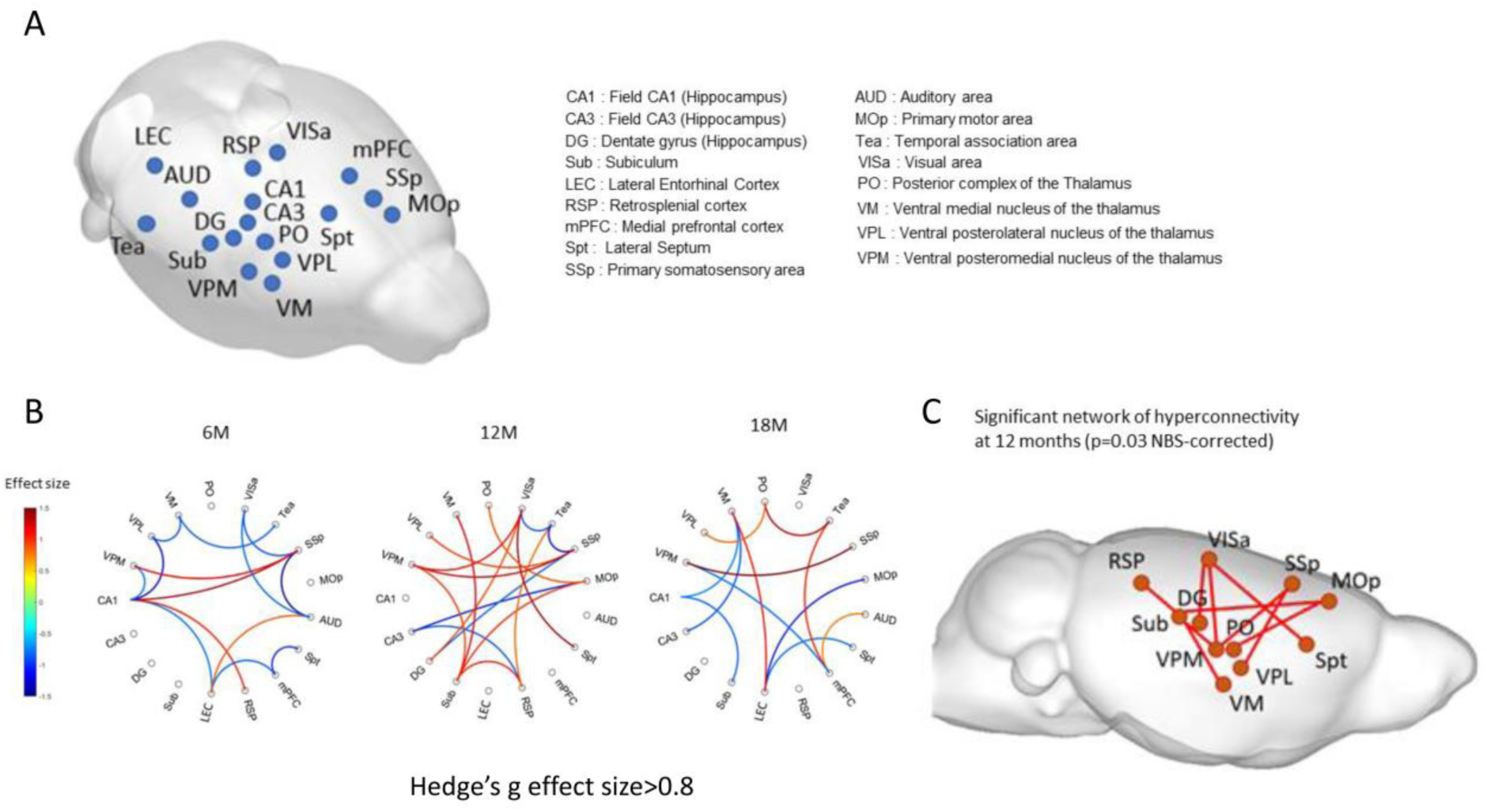
Impairment in connectivity at month 6, 12 and 18, using whole-brain rs-fMI for line 3. A. Position of the 17 ROIs used for the rs-fMRI analysis and their abbreviations. Correction for multiple comparison using NBS revealed significant networks in the hBIN1 group when compared to the WT group at each time point. B. At 6 months: identification of a network of reduced connectivity. At 12 months: network of increased connectivity. At 18 months (**d**): network of mixed reduced and increased connectivity. We represent these networks on a circular graph and in a sagittal view of the mouse brain. Hypo-connectivity is indicated in blue, hyper-connectivity in red. C. Significant network of hyperconnectivity identified at 12 months with 11 nodes (p=0.03 NBS-corrected) in a sagittal view of the mouse brain.

In contrast, at month 12, we quantified a hyper-connectivity in particular between hippocampal-entorhinal area structures and memory-related structures (Fig.4B). We identified a hyperconnectivirty network involving 11 regions. Five regions are directly related to episodic memory (DG, Sub, RSP, Lateral septum, VIsa). Mouse VIsa is equivalent to the Posterior Parietal Cortex (PPC) in primates and has been implicated in sensory and multisensory processing, navigation, motion planning, and decision-making (Lyamzin and Benucci, 2019). Furthermore, it was recently demonstrated that disease spreads from LEC to PPC that both in mouse models and LOAD patients (Kahn et al., 2014). Retrosplenial cortex is also involved in a variety of cognitive functions, including episodic memory, navigation, imagination and planning for the future (Vann et al., 2009).

Furthermore, seven of the eleven regions of the network, SUB, DG, VPM, VM, VPL, Mop and PO have been quantified with engram high index (Roy et al., 2019). RSP, VIS, SSp, Mos and Mop display direct synaptic connexions with LEC (Zingg et al., 2014). The engram approach is based on the concept that engrams are held by neuronal ensembles that are activated by learning and are reactivated to support recall (Josselyn and Tonegawa, 2020; Semon, 1920). It has been proposed that engrams for a specific memory are distributed among multiple brain regions that are functionally connected (Roy et al., 2017, 2019). Our results suggest that impacted subregions are either synaptically connected to LEC or part of a whole-brain dispersed engram complex.

At month 18, we found again a mixed hypo-connectivity and hyper-connectivity for regions linked to episodic memory (CA1, CA3, Sub, LEC, mPFC, Lateral Septum, TEA) (Fig. 4D).

Similarly, we evidenced *in vivo* an extension of the cognitive impact that was limited to impairment in object recognition component of episodic memory at month 6 (Connor and Knierim, 2017) to context pattern separation at 15 months of age (Fig. 3H) (Supplementary Figure 8). We used the protocol of context pattern separation based on two slightly distinct contexts as in (McHugh et al., 2007). Interestingly, these phenotypical changes and their spreading occurred in the absence of changes in Abeta (Supplementary Figure. 9). Using a modified MRI-DTI protocol, we were able to evidence no hypo-densities as signatures of Abeta plaques at month 18 (Supplementary Figures 10 and 11). Altogether, these results indicate a spreading from entorhinal-hippocampus region to adjacent neocortical regions and an *in vivo* impairment of both object and context episodic memory.

Mouse *Bin1* is expressed in different classes of brain cells: neurons, oligodendrocytes and microglia (Butovsky et al., 2014; Zhang et al., 2014). In the human brain, BIN1 was found with predominant white matter localization linked to oligodendrocyte expression and with overexpression in AD (De Rossi et al., 2016). A single-cell atlas of EC from individuals with AD recently revealed that BIN1 was highly upregulated in AD neurons, astrocytes, and oligodendrocytes (Grubman et al., 2019). To identify interactors of BIN1 in the different types of brain cells, we generated knockin transgenic mice expressing BIN1 with an immunoprecipitation tag either specific for neuronal isoform or ubiquitous isoforms (Supplementary Figure. 12) to identify Bin1 interactors from the whole brain (Supplementary Figures 13, 14, 15). We evidenced three types of BIN1 protein-protein interactome modules, one linked to GO Chemical synaptic transmission (P-value<2.220446e-16), one to GO Endoplasmic reticulum (P-value<2.220446e-16) and one to GO myelin sheath (p=4.56e-07) (Supplementary Table 5; Supplementary Figures 16 and 17). Interestingly, ubiquitous tagged BIN1 allowed to isolate an oligodendrocyte protein complex that includes Mog and Plp1.

We next used mass spectrometry from mouse brain synaptosomes and BIN1 immuno precipitation and identify that BIN1 interacts with proteins involved in postsynaptic complexes. We characterized two main PPI modules: Chemical synaptic transmission (with submodules: Ionotropic glutamate receptor complex and Postsynaptic density) and Endoplasmic reticulum, all of them with p-value<2.220446e-16. Sixteen genes of our DGE set were found in the submodule PPI_BIOGRID_M310: Postsynaptic density that includes 120 distinct proteins (Supplementary Table 6; Supplementary Figure 18). Furthermore, from differential gene expression data of laser-assisted microdissection of *hBIN1* hippocampal subregions, we evidenced genes specifically expressed in distinct glia lineages, astrocytes, oligodendrocytes and microglia with an enrichment (Pval<0.0055) in deregulated microglia genes (Zeisel et al., 2015) (Supplementary Tables 7,8). Altogether, these results indicate that deregulation of BIN1 can impact astrocytes, oligodendrocytes, microglia and neurons. In order to detect a possible signature of neuroinflammation, we used small-molecule radio-ligands specific for the translocator protein (TSPO) (18 kDa) that has been used extensively to detect microglial activation and neuroinflammation in both preclinical and clinical studies (Chen and Guilarte, 2008; Owen and Matthews, 2011). We did not detect any significant changes in the hippocampus-entorhinal cortex region in *hBIN1* mice compared to control at month 15, indicating that possible changes are under the sensitivity of this approach (James et al., 2015) (Supplementary Figure 19). Altogether, these results indicate early changes in the different types of brain cells in our hBIN1 model.

We next studied localization of BIN1 at neuronal synapses using an improved direct Stochastic Optical Resolution Microscopy (dSTORM) (Bourg et al., 2015) (Fig. 5A, B, C, D). BIN1 was identified as part of PSD and of synaptosomal proteins. We found BIN1 protein is present in both pre and post-synapse with a two-fold increase in the number of molecules in the post compartment as compared to presynaptic compartment (Fig. 5c, d). These data are in full agreement with our mass spectrometry data from synaptosomes (Supplementary Table 6; Supplementary Fig. 13) and previous proteomics studies on PSD fraction (Collins et al., 2006). We further analyzed molecular mechanisms that may generate spine impairments of DG neurons as evidenced by diI biolistic injection of *hBIN1* mice hippocampal slices at 3 months (Fig.3 A, B). From our yeast two-hybrid studies and Biogrid data, we found that BIN1 interacted with DLG1 and with the Rho/Rac1 Guanine nucleotide exchange factors (GEF), KALRN and TIAM1, interactors of RAC1 and involved in dendritic spine and/or neurite growth (Goto et al., 2011; Luo et al., 1996; Penzes and Remmers, 2012; Tashiro et al., 2000; Um et al., 2014) (Fig. 5E). We validated BIN1-TIAM1 and TIAM1-DLG1 interactions by proximity ligation assay, demonstrating the postsynaptic location of DLG1, TIAM1 and BIN1 (Fig. 5F). We next tested the hypothesis that overexpression of BIN1 at the post-synapse can induce a long-lasting activation of Rac1. This activation of Rac1 can induce changes in actin polymerization of dendritic spines able to modify both the morphology of spines (Caroni et al., 2012; Hayashi-Takagi et al., 2010) and the long-term potentiation of synapses (Rust et al., 2010). Rac1 is active when bound to GTP and is inactive when bound to GDP. Rac1 activation was detected by increased GTP-bound Rac1 and its specific binding to the p21-binding domain (PBD) of p21-activated protein kinase 1 (Pak1). Here, we found that GST-Pak1-PBD levels were doubled in *hBIN1* hippocampus compared to control (Fig. 5G), suggesting that overexpression of BIN1 can impact structural plasticity of dendritic spines of DG neurons via changes in Rac1 activity.

**Figure 5.**
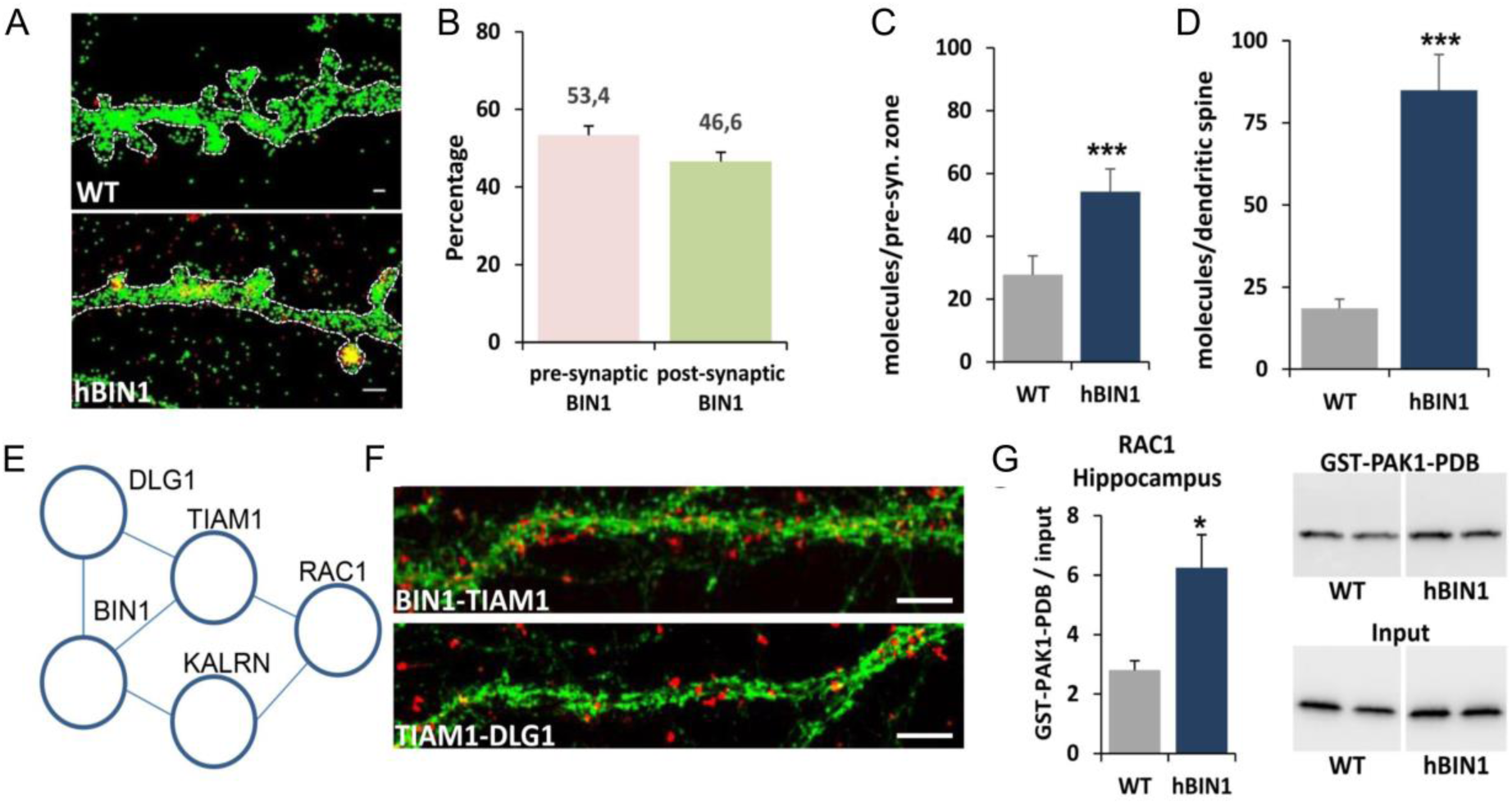
Analysis of specific synaptic defects in hBIN1 mice. BIN1 is expressed in postsynaptic dendritic spines. d-STORM acquisition of BIN1 (red) and Phalloïdin (green) on DIV21 WT and hBIN1 hippocampal neurons. **A.** Percentage of BIN1 molecules detected on pre-synaptic and post-synaptic compartment on Wild-type neurons. Scale bar=0,5µm. **B.** Quantification of BIN1 molecules detected using d-STORM in pre-synaptic compartment in WT (n=118) and in hBIN1 (n=170) neurons from n= 3 WT and n=4 hBIN1 embryos. *** p < 0.0005. **C.** Quantification of BIN1 molecules detected using d-STORM in post-synaptic compartment in WT (n=118) and in hBIN1 (n=170) neurons from n= 3 WT and n=4 hBIN1 embryos. *** p < 0.0005. **D.** Cytoscape network of BIN1 interactions obtained by yeast-two-hybrid tests (TIAM1, KALRN, DLG1) and from Biogrid (RAC1). **E.** *In situ* PLA (red) using anti-BIN1 and anti-TIAM1 or anti-TIAM1 and anti-DLG1 antibodies co-labelled with actin (Phalloïdin, green) on WT mice primary hippocampal neurons 21DIV. Scale bars=5μm. **F.** Western blot analyses of GTP-Rac1 pull-down bound with GST-PAK1-PDB protein fusion and total Rac1 from WT (n=8) and hBIN1 (n=4) hippocampi, showing an increase of Rac1 active forms (*p<0,05).

## Discussion

Our new finding is that the LOAD genetics risk factor BIN1 overexpression is sufficient to induce deregulated repertoires of co-expression genes that are common to mouse hippocampal subregions of *hBIN1* mice at 7 weeks and to human post-mortem brain samples from Alzheimer patients. This common deregulation validates this LOAD genetic risk factor-based mouse model.

The second new finding is the first demonstration, to the best of our knowledge, that a LOAD genetics risk factor can alone induce structural and functional changes in the absence of any modifications of Aβ1-42 peptides. The structural changes are centered on the LEC and its direct synaptically-connected targets that are known to be vulnerable regions in early phases of LOAD (Braak and Braak, 1995; Braak et al., 1993). Recent studies evidenced that it is the lateral, rather than the medial, entorhinal cortex that is most susceptible to tau pathology early in AD, with spreading to other cortical sites (Khan et al., 2014a). From these results, we propose that multiple factors can impact such vulnerable sub-regions, including tau, APP and LOAD genetics risk factors such as BIN1, leading to early EC-DG phenotypes with distinct properties.

Neurodegeneration can be defined as a progressive loss of neurons or their processes (axons and dendrites) with a corresponding progressive impairment in neuronal function (Jack et al., 2013). Here, we evidenced neurodegeneration spreading of both structural tract alterations and connectivity using whole-brain MRI-DTI and rs-fMRI, with aging. Such phenotypical progression occurs in the absence of any Abeta aggregates. This suggests that progression of phenotype via neuronal network connections such as entorhinal cortex-> temporal association cortex>somatic sensory-motor subregions (see Fig.7 in (Zingg et al., 2014)) can be linked to trans-synaptic propagation of the disease involving molecular mechanisms that requires further studies using this mouse preclinical model. This *hBIN1* mouse model can be instrumental to study spreading of neurodegeneration in the absence of either Abeta peptides changes.

We found repertoires of co-deregulated genes that display specific expression in subsets of glial cells, oligodendrocytes, astrocytes and microglia. All these three compartments are known to be involved in synaptic homeostasis and to change in LOAD (Chung et al., 2016; Hong et al., 2016; Keren-Shaul et al., 2017; McKenzie et al., 2017; Zhang et al., 2013). Experience-dependent enhancement of myelination in the mature cortex may accelerate information transfer in these circuits and strengthen the ability of axons to sustain activity by providing additional metabolic support (Hughes et al., 2018; McKenzie et al., 2017). Combined modifications of gene expression in these three types of glial cells can generate early structural tract alterations that we report here. Furthermore, these non-neuronal events are expected to impact the normal function of BIN1 at the post-synapse. Further work will be needed to identify the spatial and temporal sequence of these events. We propose that the combination of glial defects in a specific sub-region contribute with synaptic BIN1 overexpression to a dysfunction restricted to synapses of EC-hippocampal pathways. The synaptic dysfunction in vulnerable subregions such as EC-Dentate Gyrus performant path can lead to an abnormal structure of dendritic spines related to a deregulated Rac1 activity. This Rac1 pathway can be a candidate for further manipulation in order to evaluate potential rescue of cognitive defects.

Altogether, our findings underscore the potential of preclinical mouse models based on LOAD genetic risk factor to identify co-regulated gene repertoires as novel biomarkers able to be manipulated in the silent phase of AD.

## Material & Methods

### Differential gene expression analysis

We extracted the gene expression and clinical information from BM36 of a human AD brain study (syn3157743). We grouped the subjects into 3 categories: nondemented (CDR=0, n=31), mild (CDR=0.5 or 1, n=54) and severe (CDR=2 and above, n=116) and perform DEG analysis using Limma in R 3.3.2 (Ritchie et al., 2015).

### hBIN1 hippocampus subregions Laser Microdissection and microarrays assay

Snap frozen brains from seven-weeks-old male mice were cut using a cryostat (Cryostat HM550 MM, Thermo Scientific) in sagittal 40 µm slices from lateral 3.00 to lateral 2.52 mm. Slices were then colored with Hematoxylin-Eosin and dehydrated using 50%, 70% and 100 % Ethanol solutions. The dorsal and ventral structures (CA1/2, CA3, Dentate Gyrus) of the hippocampus were microdissected using the PalmRoboPro software and the PALM® Microbeam (Carl Zeiss MicroImaging GmbH) Laser microdissecter. Tissues were collected in 30 µl RNA Extraction Buffer (Arcturus Picopure RNA isolation kit Applied Biosystems) and RNA was extracted according to the Arcturus Picopure kit instructions. Microarray assays were performed by IGBMC, Plateforme Biopuces et Séquençage, Strasbourg, France, using Affymetrix technology with GeneChip® Mouse Gene 2.0. Analysis was made using the FCROS method (fold change rank ordering statistics) as previously described (Dembélé and Kastner, 2014). The expression was considered as significantly different between WT and transgenic mice when p-value was under the 0.02 treshold.

### Construction of *BIN1*-centered correlation network from human AD studies

We constructed a *BIN1*-centered correlation network in AD based on eight gene expression datasets from three independent human AD cohorts including the Harvard Brain Bank[1], the ROSMAP study (doi:10.7303/syn3219045) and Mount Sinai Brain Bank (https://www.synapse.org/#!Synapse:syn3157743). The correlation between Bin1 and all other genes were calculated on gene level in each individual datasets. Genes with multiple probes were represented by the probe with the largest variation. The gene correlation with a BH-corrected p value <0.05 was considered significant. Fisher’s Exact Test (FET) was performed to test the enrichment between DEG signatures from *hBIN1* transgenic mice and *BIN1*-correlated genes that were consensus across multiple human datasets.

### hBIN1 mouse line generation and validation

The humanized hBIN1mouse lines were established at the MCI/ICS (Mouse Clinical Institute - Institut Clinique de la Souris, Illkirch, France; http://www-mci.u-strasbg.fr). A BAC (RP11-437K23 - RPCI human BAC library 11) originally generated by BACPAC Resource Center (BPRC) at Children’s Hospital Oakland Research Institute in Oakland (California) was purchased. A maxiprep (NucleoBond® BAC 100; Macherey Nagel) was prepared. A careful restriction digest was performed in order to verify the integrity of the BAC and the sequence of all coding exons was confirmed by PCR and Sanger sequencing. The BAC prep was microinjected in the pronuclei of C57Bl/6N fertilized oocytes. Eleven founders were obtained and bred for germ line transmission.

The number of BAC copies for each established sublines was estimated by qPCR (LightCycler® 480 SYBR Green I Master) on ears DNA from 2 transgenic animals compared to one wt reference: murin DNA copies was measured after sequences amplication targeted on intron5/exon6 junction (primers sequences Fw1: gccccactgatctctcctc / Rv1: ggcactgtcatagtccacca). BAC human DNA copies was quantified by amplication of exon 2 region (primers sequences Fw2: tccccttctctcttggcttc / Rv2: ctgctcatccttggtctcatc) and final region of intron 19 (primers sequences Fw3: cttcccagcaggttggagtgg / Rv3: accaggtcgcacagggatg). Then, we looked at the mRNA expression by RT-qPCR: reverse transcription was done on 1 µg of RNA with the QuantiTect® Reverse Transcription from qiagen. cDNA was produced from liver, brain and quadriceps extracts of one wt sample and three transgenic mice. Two sets of primers were designed aiming at exon 20 to distinguish murin specific cDNA (Fw4: attttacagagcgggtgcag / Rv4: ctttgggggaaaggttcttc) and human version (Fw5: cactgagagggtcccatgac/ Rv5: cacacatttttcgggaggag). On a second step, we designed two other cuples of primers to separately amplify the neuronal isoforms (1-7), hybriding in exon 13 (Fw6: ggacacgtttgtccctgag / Rv6: ccttcacagggctcgtca) and another located in the muscle specific isoform (8) located in exon 11 (Fw7: acgggagcaacaccttca / Rv7: gccgcgaaaacagtttactt)

Protein overexpression was validated by western blot quantification. Hippocampus and muscle samples were collected from 3 mice of each genotypes wt and Tg(*BIN1)* at 3 months of age, and 4 samples of each genotypes at 15 months. Tissues were crushed with Precellys system (Lysing Kit CK14) in RIPA lysis buffer (santa cruz sc-24948). The protein concentration was determined by Bradford dosage with Pierce BCA protein assay kit (Thermo scientific #23225). Ten microgram of total protein of hippocampus and muscle extracts were electrophoretically separated in SDS–polyacrylamide gels (10%) and then transferred to nitrocellulose membrane (120V) during 1h30 on ice. Non-specific binding sites were blocked with 5% skim milk powder in Tween Tris buffer saline 1 h at room temperature. Immunoblotting was carried out with Bin1 pan isoforms primary antibody (mouse anti-Bin1 1/5000, sigma B9428) and followed by secondary conjugated with horseradish peroxidase (goat anti mouse dako P0448). The immunoreactions were visualized by ECL chemiluminescence system (Pierce ECL western blotting substrate – thermo scientific); Epifluorescence were captured with Amersham Imager 600. Bands were detected at 85 and 90 kDa in the brain, and 55 and 64 kDa in the muscle; intensities were quantified with ImageJ.

### Ethical Statement and mouse breeding

Mice were handled with the agreement of the local ethical committee and in accordance with the European Council Guidelines for the Care and Use of Laboratory animals ETS 123. Dr Yann HERAULT was granted the accreditation 67-369 to perform the reported experiments. All animals were treated in compliance with animal welfare policies from the French Ministry of Agriculture. Our project was granted permission by the French Ministry of Agriculture (law 87 848) under the accreditation 67–369. We had the approval from our local ethical committee to realize our experiments through the accreditation number 2014-056. For all these tests, mice had free access to water and food (D04 chow diet, Safe, Augy, France). The light cycle was controlled as 12 h light and 12 h dark (lights on at 7 am) and the temperature was maintained at 23±1 °C.

Behavioral studies were performed between 3 and 15 months of age. Mice were transferred from the animal housing facility to the phenotyping area at the age of 8-10 weeks. On testing days, animals were transferred to experimental room antechambers 30 min before the start of the experiment. All experiments were performed between 9:00 AM and 4:00 PM. Mice had at least 3 days of rest between 2 consecutive tests. To study the behavior of animals, we crossed transgenic hemizygotes mices (Tg(*BIN1*)/0) with wild-type C57BL/6J animal to generate 3 successive cohorts of males that were studied at different ages. The cohort n°1 with 9 Tg(*BIN1*)/0 and 10 wildtype (WT) littermates was studied at 3, 6, 9, 12 and 15 months. To confirm the phenotypes observed in this cohort, and to avoid the effect of cumulative tests, we produced 2 other cohorts: one was tested at 3 and 6 months (cohort n°2), and the other (cohort n°3) at 9 and 12 months. In the second cohort were 13 Tg(*BIN1*)/0 and 15 wt littermates and in the third cohort 11 Tg(*BIN1*)/0 mice and 12 wt littermates. Tests were run in the following order: open field, novel object recognition, fear conditioning in a period of 3 weeks that are described in the supplementary information.

### MRI experiments

All MRI experiments were performed on a 17.2T horizontal bore animal MRI system (Bruker BioSpin, Ettlingen, Germany) equipped with gradients of 1000mT/m-maximum strength using a dedicated mouse brain quadrature birdcage coil (Rapid Biomedical GmbH, Rimpar, Germany).

All animal experimental procedures were performed in accordance with the EU Directive 2010/63/EU for care and use of laboratory animals and approved by the Comité d’Ethique en Experimentation Animale (CETEA), de la la Direction de la Recherche Fondamentale (DRF) du Commissariat à l’Energie Atomique et aux Energies Alternatives (approval number: APAFIS#8462-20170109l5542l22 v2).

### Ex vivo DTI study

#### Subjects

In total, we scanned 55 ex-vivo mouse brains: 26 at 3 months and 29 at 6 months. The 3-month-old group was divided into 3 subgroups: 10 wild-type controls (WT), 6 hBIN1 U154.16.16 (line 1) and 10 hBIN1 U154.45.12 (line 2). The 6-month-old group was divided into 3 subgroups: 10 wild-type controls (WT), 10 hBIN1 U154.45.12 (line 2), and 9 hBIN1 U154.20.8 (line 3).

#### Tissue preparation

After intracardiac perfusion (4% paraformaldehyde + Gd-DOTA), brains (within the skull) were collected and chemically fixed by immersion in 4% paraformaldehyde + Gd-DOTA for 24 hours. After fixation, the brains were placed in PBS + Gd-DOTA for storage. For imaging the samples were placed in fluorinert (FC40, Sigma-Aldrich, L’Isle d’Abeau Chesnes, France).

#### Diffusion

*MRI acquisitions* A 3-D diffusion-weighted Pulsed Gradient Spin Echo Echoplanar (EPI) sequence was implemented with the following parameters: 13-segments, TR/TE=250/24.5ms, diffusion gradients duration/spacing δ/Δ = 5.0/12.3ms, b-value 1500s/mm^2^ with 25 directions and 17 b=0s/mm^2^ reference images, isotropic resolution of 100µm, matrix size=192×152×152, total acquisition time of 4h. A 3D T2-weighted TurboRARE acquisition with 60µm isotropic resolution was also performed for registration purposes.

#### Data analysis

Data preprocessing was carried out using the Ginkgo toolbox available at https://framagit.org/cpoupon/gkg ([CSL STYLE ERROR: reference with no printed form.]), including the registration of each subject anatomical image to its diffusion space, correction of rician noise using a non-local means filtering algorithm and correction for eddy currents. A 3D mouse brain atlas (Turone mouse brain atlas, (André Barrière et al., 2020) was then normalized to each subject space. For analysis, we selected in total 16 ROIs, including 10 global ROIs spanning the entire brain and 6 hippocampal-entorhinal ROIs.

The Ginkgo toolbox was used to compute the DTI model and derive the FA and MD quantitative maps. FA and MD values were averaged over all voxels of each bilateral ROI of each mouse, then averaged for each ROI over all mice of the same group.

#### Statistical analysis

To compare the 3 subgroups at each age, we used Tukey’s HSD test, a single-step multiple comparison procedure which corrects for the family-wise error-rate. Significance level was set to p=0.05.

### In vivo resting-state fMRI study

#### Acquisitions

Two groups of mice, 11 controls and 10 hBIN1, were scanned at 3 time-points: 6, 12 and 18 months of age. Mice were anesthetized with isoflurane 0.5% + medetomidine (0.05 mg/kg/h, s.c.). After a localizer scan and local shimming, an anatomical image was acquired using a RARE sequence with the following parameters: 12 slices, slice thickness=400 µm, resolution 100×100 µm. The rs-fMRI data were acquired with a GE-EPI sequence with the following parameters: TE=10 ms, TR=1.5s, 12 slices, slice thickness = 400 µm, 300 repetitions, in-plane resolution 200×200 µm. Total scanning time was 45 minutes per mouse.

#### Preprocessing of images

SPM12 software *(Wellcome Trust Center for Neuroimaging, UK)* was used for preprocessing, including slice timing correction, motion correction by realignment, coregistration of the functional image to a structural brain image, spatial normalization, and smoothing with a gaussian filter (FWHM = 0.2 mm). Functional data denoising was applied with CONN toolbox (Whitfield-Gabrieli and Nieto-Castanon, 2012), including regression of signal due to head motion with 12 parameters (6 realignment parameters + 6 derivatives), aCompCor (Behzadi et al., 2007) regressors for CSF and white matter (5 components each), scrubbing based on framewise displacement (Power et al., 2012), bandpass filtering (0.01 to 0.1 Hz) and linear detrending.

*Regions of interest (ROIs)* were extracted from an atlas (Turone Mouse Brain Template and Atlas).

#### Resting-state functional connectivity

Estimation of ROI-to-ROI correlations for each mouse was done with CONN toolbox. The correlation coefficient was computed between preprocessed and regionally averaged (17 regions) BOLD signals and converted to Z-scores using Fisher’s r-to-z transformation, to yield a connectivity matrix for each mouse. Regions of interest (ROIs) were originally created using the Allen Brain Atlas (Oh et al., 2014) and registered to a mouse brain MRI template to create a labeled atlas.

#### Network-based statistics

The NBS approach *(A. Zalesky et al. 2010)* was used to identify subnetworks in which correlation strength (Z-scores) significantly differed between the WT group and the hBIN1 group at each timepoint. This procedure separates sets of highly interconnected regions (subnetworks), instead of paired regions, while controlling for the family-wise error rate by permutation testing. The NBS was performed as follows: for each of the 17 × 16/2 = 136 unique pairs of ROIs (functional connections), a two-sample t-test was used to independently test whether group-averaged functional connectivity differed between the two groups. This yielded a 17 × 17 matrix of t-statistic values. Then, Hedge’s g (effect size) of group difference in each subnetwork was calculated from t-values. The subnetwork with an effect size exceeding 0.8 (large effect) was used for further NBS analysis. Specifically, 5000 permutations were generated in which group labels were randomly permuted. Subnetworks deemed to be significant with permutation testing (p < 0.05, familywise error-corrected) were visualized. The NBS analysis were performed with the NBS Connectome software (www.nitrc.org/projects/nbs).

### Tissue Preparation and Sectioning

Mice were deeply anesthetized with a Ketamin/Xylazine mix (10 µl/g body weight) and perfused transcardiacally. For DiI assays, perfusions were made with a solution containing 2% paraformaldehyde. Brains were dissected out and fixed 1h in the same perfusion solution at 4°C. Coronal sections (70 µm thick) were cut with a vibratome and kept at 4°C in a PBS1X Azide Na 0.01% solution until the experiment.

### Immunohistochemistry

Brain sections were rinsed three times in PBS1X and permeabilized in a PBS1X 0.2% Triton solution (30 minutes, room temperature). DNA was denatured using a PBS1X HCl 2N solution (30 minutes, 37°C) and sections were incubated with rat anti-BrdU antibody (see Supplementary table) overnight at 4°C. Sections were then incubated with anti-rat biotinylated secondary antibody (2h, room temperature) and processed with avidin-biotin-peroxydase complex (ABC Elite Kit, Vector Laboratories) for 30 minutes. Finally, peroxidase detection was conducted with 3,3diaminobenzidine-tetrahydrochloride as chromogen with H2O2 (Kit SK-4100, Vector Laboratories). Sections were then mounted in Dako mounting medium.

### DiI Assays

DiI cristals were spread on brain sections using the Helios Gene Gun System (Biorad). Sections were incubated 3h with DiI and fixed in PFA 4% for 30 min at room temperature. After three times in PBS1X, sections were mounted in mowiol. Acquisitions were made using confocal microscope (Leica, SP5 from PICPEN imagery platform Centre de Psychiatrie et Neuroscience) at 40× magnification, zoom 4, z-stack 0.21 µm. Spines were analyzed with Neurolucida software on 3D reconstructions. The density of each type of spine (mushroom, stubby, filopodia) was quantified for one dendrite per neuron.

### Dendrite organization analysis

To evaluate the effect of BIN1 overexpression on dendritic arborisation a Golgi-Cox staining method was applied on 4 males TgBIN1 and 4 males WT littermate at two months. After dissection brains were placed in 12 ml Golgi–Cox solution (potassium dichromate 5%, mercuric chloride 5% and potassium chromate 5%), where they were stored in the dark for 40 days at RT and solutions were refreshed after 2 days. Following this incubation, brains were then transferred to a 30 % sucrose solution for at least 3 days and coronal sections (200 µm) containing hippocampus were obtained using a vibratome (Leica vt 1000S). Sections were collected into 24-wells plate in sucrose 6% for one day stored at 4°C. Tissue slices were developed in a series of solutions (Ammonium hydroxide, Kodak Fix solution, 50% ethanol, 70% EtOH, 95 %EtOH, 100% EtOH and chloroform, xylene and alcohol mix) and mounted on slides with pertex, and coverslipped (Levine et al., 2013).

Three dimensional tracing of neurons, in dentate gyrus (DG), was carried out using a 63X objective on a Leica TCS SP5 CFS equipped with a white light laser. To detect Golgi-Cox material, as previously described by (Spiga et al., 2011). The microscope was automatically set up in reflection mode by using a 488 nm. Subsequent analyses of dendritic extent were performed with simple neurite tracer from Fiji©. Briefly, criteria for selection included: (a) full impregnation of the neurons along the entire length of the dendritic tree; (b) dendrites without significant truncation of branches and (c) relative isolation from neighboring impregnated neurons, astrocytes or blood vessels. Five to sixteen granular cells were reconstructed per animal, resulting in the analysis of thirty to forty-four neurons per experimental group. Sholl analysis was performed by using Fiji software and in accordance with Ferreira et al. plugin for 3 dimension sholl analysis (Ferreira et al., 2014). Sholl analysis data were compared across groups using repeated measures ANOVA followed by Bonferroni post-test analysis. Significance was set at p < 0.05.

### Ex vivo electrophysiology

Electrophysiological experiments were performed in WT (n>10) and BIN1-HET (n>10) mice. After decapitation of a mouse under deep anaesthesia, the brain was rapidly removed from the skull and placed in a chilled (0-3°C) artificial cerebrospinal fluid (ACSF) containing (mM) NaCl 124, KCl 3.5, MgSO4 1.5, CaCl2 2.5, NaHCO3 26.2, NaH2PO4 1.2, glucose 11. Transverse slices (300-400 µm thick) were cut using a vibratome and placed in a holding chamber (at 27°C) containing the ACSF solution, at least one hour before recording. Each slice was individually transferred to a submersion-type recording chamber and submerged with ACSF continuously superfused and equilibrated with 95% O2, 5% CO2. Recordings of presynaptic fiber volleys (PFV) and field excitatory postsynaptic potentials (fEPSPs), mostly resulting from the activation of AMPA receptors, construction of Input/Output (I/O) curves assessing the responsiveness of AMPA/Kainate glutamate receptor (A/K), and measure of paired-pulse facilitation (PPF) have been previously detailed (Potier et al., 2010). Long-term potentiation (LTP) was produced with theta-burst stimulation, consisting of five trains of four 100 Hz pulses each, separated by 200 ms and delivered at the test intensity (repeated four times with an interburst interval of 10 s). Another type of stronger LTP was also produced by application of a 3×100 Hz stimulation spaced of 20 sec. Long term depression (LTD) was induced by applying a low frequency stimulation at 2 Hz (1200 pulses for 10 min).

Experiments were done in the CA1 area following stimulation of stratum radiatum, or in the dentate gyrus (DG). For DG experiments, stimulation and recording electrodes were both placed in the middle one-third of the DG molecular layer, 500 µM apart and bicuculline methiodide (50 µM, Sigma-Aldrich, Saint Quentin Fallavier, France) was added to the perfusing medium throughout the LTP experiment.

Measurement of the biophysical properties of the glutamate-dependent or GABA-dependent tonic currents, as well as miniature spontaneous activity dependent on these neurotransmitters was performed in hippocampal slices from BIN1 mice, compared to wild-type siblings. Whole-cell patch-clamp recordings of CA1 pyramidal neurones or dentate gyrus granular cells were performed at room temperature, using borosilicate patch pipettes (5 MΩ) filled with CsCH4O3S 140 mM, CsCl 6 mM, MgCl2 2mM, HEPES 10 mM, EGTA 1,1 mM, QX-314 5 mM, ATP 4 mM, (pH 7.3; 290 mosm).

Transmembrane currents were acquired and filtered by an amplifier (AxoPatch 1-D, Axon Instruments), stored on a computer and digitalized using the WinLTP software (Anderson and Collingridge, 2001) for on-line and off-line analysis.

Miniature spontaneous excitatory post-synaptic currents (mEPSCs) were recorded in the continuous mode at hp = −60 mV, in the presence of TTX (1µM) and bicuculline (10 µM). The glutamate-dependent tonic current was recorded at a holding potential of + 40 mV, in the presence of TTX (1 µM), NBQX (10 µM), and bicuculline (10 µM) to isolate the NMDA component of the holding current (hc). After 3-5 minutes recording of a stable control hc, APV (50µM) was applied to the superfusion medium. The hc dropped to a new stable value under the effect of APV, and the difference between hc (control) with hc (APV) determined the amplitude of the tonic Glu current.

The GABA-dependent miniature inhibitory post-synaptic currents (mIPSCs) were recorded at +10 mV in the presence of TTX (1µM), NBQX (10µM) and APV (50 µM). In this configuration, the addition of bicuculline (10µM) fully abolished the mIPSCs and induced a drop of the hc to a new stable value. The difference between hc (control) and hc (bicu) determined the amplitude of the tonic GABA current. Frequence and amplitude measures of mEPSCs and mIPSCs were performed using Spike 2 (CED, Cambridge).

### Behavioral Procedures

#### Novel Object Recognition paradigm

To assess recognition memory, we used the Novel Object Recognition task. On day 1 (D1), habituation session was carried out in a 55 cm diameter white round box and mouse activity was recorded with a video tracking system (Ethovision, Noldus, France) during 30 min. After each mouse trial, the arena was thoroughly cleaned to minimize olfactory cues. On day 2 (D2), animals were presented with a pair of identical objects (marble or dice) until they had explored the objects more than 3 seconds, in a maximum period of 10 min starting from the first sniffing behavior. The exploration is considered as any deliberate investigation towards each object in a distance more or less than 2 cm, or when touching with the nose. Mice exploring under this threshold were excluded. In the second trial (test phase, 1h later), one of the familiar objects was changed for a new one, and the animals were left in the arena during 10 min. The same threshold of exclusion was applied after this retention phase. To control for odour cues, the OF arena and the objects were thoroughly cleaned with 50% ethanol, dried, and ventilated for a few minutes between mice. Memory was operationally defined by the time spent on each object and the discrimination index (DI): time spent investigating the novel object minus the familiar one (DI = (Novel Object Exploration Time/Total Exploration Time) – (Familiar Object Exploration Time/Total Exploration Time) X 100).

#### Morris water maze spatial memory

We chose for the task the one position learning protocol adapted from Morris (Morris, 1984). The water maze was a circular pool (150-cm diameter, 60-cm height) filled to a depth of 40 cm with water maintained at 20°C–22°C, made opaque using a white aqueous emulsion (Acusol OP 301 opacifier) and split into 4 virtual quadrants: South-East (SE), North-West (NW), North-East (NE), South-West (SW). The escape platform (PTF), made of rough plastic, was submerged 1 cm below the water’s surface. This experiment was performed to study reference memory through a spatial search strategy that involved finding the hidden platform. The spatial memory session consisted of a 6-day (D1 to D6) learning phase with four 90-second trials per day. Each trial started with mice facing the interior wall of the pool and ended when they climbed onto the platform located on the SE quadrant or after a maximum searching time of 90 sec. The starting position was changed pseudo-randomly between trials. Mice were left dried out under heating lamp during 10-15 min inter-trial intervals. On the seventh day, mice were given the 60-sec probe test in which the platform had been removed. On day 9 (C1), we performed a cued version test; this final day was based on 4 trials of 60s where the PTF, located in West, is labelled by a flag. This special test allowed to distinguish an impairment in spatial learning versus ocular defects. The latency to find the platform, distance traveled in each quadrant (NW, NE, SW, SE) and the average speed were recorded to quantify the time spent in the target quadrant. The swimming path was recorded and analyzed with the Ethovision software from Noldus®.Please note that 10 wt and 9 Tg mice realized this paradigm but 2 wt were excluded since they didn’t swim at each trial from D2.

#### Pattern Separation analysis based on contextual Fear Discrimination Task

The procedure was adapted from (McHugh et al., 2007) with Polymodal operant chambers (Coulbourn Instruments, Allentown, PA) used for this experiment. We created two different context environments: **Context A** in chamber A was defined as a room lit with overhead lighting at 50 lux and containing two conditioning chambers. The 18.5 × 18 × 21.5 cm chamber had plexiglass wall in front and back and aluminum side walls. The chamber floor consisted of a grid composed of 16 stainless steel rods connected via a cable harness to a shock generator. The chambers were cleaned between mice with 5% sodium hydroxide; a pan coated of benzaldehyde in 100% alcohol (0.25% concentration) was placed beneath the chambers during the experiment to provide an olfactory cue. **Context B** in chamber B had crucial modification: the light overhead location was changed, we masked the wall with black Plexiglas squares and we change the ceiling motif. As in context A, the floor of each chamber consisted of 16 stainless steel rods which were wired to a shock generator and scrambler. This context was cleaned and scented with a 1% acetic acid solution. The room was lit with a 30-W red overhead light. Each day, the animals were transported in their home tubs to a room adjacent to the experimental room. They were left undisturbed for at least 20 min.

On days 1-3 the mice were carried to the A-context conditioning room and placed into the conditioning chambers. After 192sec, they received a single footshock (2 sec; 0.65mA) and were removed from the chambers 1min following footshock termination. Across the subsequent two consecutive days (days 4 and 5), mice were placed into the A-context and B-context conditioning chambers in separate tests counterbalanced order). Each test consisted of an 8min exposure to the chamber without the delivery of footshock. On days 6 through 12, mice were exposed to both context A and B conditioning chambers daily. The order of exposure on each day followed a BAABABBABAAB design. Across the entire discrimination phase, all animals received a single footshock during each Chamber A exposure and never received footshock during Chamber B exposures. The dependent measure employed was freezing behavior; the general activity of the animals was recorded through the infrared cell placed at the ceiling of the chambers and was recorded directly on a PC computer using Graphic State (Coulbourn). Score of each mouse represented freezing or not freezing state every 2 seconds. These scores were then converted into a percentage of observations spent freezing. For the 12 days, we evaluated the percentage of freezing only on the 3 first minutes after mice were placed in the chamber (192s for conditioning days).

#### Yeast Two-Hybrid Analysis

Yeast two-hybrid screening was performed by Hybrigenics Services, S.A.S., Paris, France (http://www.hybrigenics-services.com).

The coding sequence for mouse amphiphysin 2 (aa 1-410; GenBank accession number gi: 134053916) was PCR-amplified and cloned into pB27 as a C-terminal fusion to LexA (N-LexA-Amph-C). The construct was checked by sequencing the entire insert and used as a bait to screen a random-primed Mouse adult Brain cDNA library constructed into pP6. pB27 and pP6 derive from the original pBTM116 (Vojtek and Hollenberg, 1995) and pGADGH (Bartel et al., 1993) plasmids, respectively.

77 million clones (7-fold the complexity of the library) were screened using a mating approach with YHGX13 (Y187 ade2-101::loxP-kanMX-loxP, matα) strains and L40ΔGal4 (mata) yeast strains as previously described (Fromont-Racine et al., 1997). 131 His+ colonies were selected on a medium lacking tryptophan, leucine and histidine. The prey fragments of the positive clones were amplified by PCR and sequenced at their 5’ and 3’ junctions. The resulting sequences were used to identify the corresponding interacting proteins in the GenBank database (NCBI) using a fully automated procedure. A confidence score (PBS, for Predicted Biological Score) was attributed to each interaction as previously described (Formstecher et al., 2005).

##### Further description of the confidence score

The PBS relies on two different levels of analysis. Firstly, a local score takes into account the redundancy and independency of prey fragments, as well as the distribution of reading frames and stop codons in overlapping fragments. Secondly, a global score takes into account the interactions found in all the screens performed at Hybrigenics using the same library. This global score represents the probability of an interaction being nonspecific. For practical use, the scores were divided into four categories, from A (highest confidence) to D (lowest confidence). A fifth category (E) specifically flags interactions involving highly connected prey domains previously found several times in screens performed on libraries derived from the same organism. Finally, several of these highly connected domains have been confirmed as false-positives of the technique and are now tagged as F. The PBS scores have been shown to positively correlate with the biological significance of interactions.

#### Autoradiography with [^18^F] DPA-714

For autoradiographic studies 9-10 months old WT and hBIN1 transgenic mice were perfused with saline. Brain tissue was frozen immediately and kept on −80°C until use. [^18^F]DPA-714 (68GBq/µmol; 3nM) autoradiographic studies were performed using 20 µm axial brain sections. [^18^F]DPA-714 radiosynthesis was prepared as previously described (Pottier et al., 2017). Specific binding was assessed using an excess (20 µM) of unlabeled DPA-714. Briefly, sections were incubated for 20 min in Tris Buffer (50mM TRIZMA preset crystals purchased from Sigma-Aldrich) adjusted to pH 7.4 with NaCl. The unbound excess ligands were removed by two wash cycles (2 min each one) in cold buffer and then a final rinse in cold deionized water. Sections were then placed in direct contact with a Phosphor-Imager screen (Molecular Dynamics, Sunnyvale, CA) and exposed overnight. Autoradiograms were analyzed using ImageJ (NIH) software. For quantification, we analyzed at least six sections per animal (n=3/genotype). For each section, a region was drawn within the striatum (left & right) and another one including the dentate gyrus and the medial entorhinal cortex (left & right) between bregma −4.44 and −6.24. The mean grey values measured for each section were normalized to the mean grey value of the corresponding striatum.

#### Primary cell culture

E17.5 hippocampal neurons were enzymatically dissociated (0.25% trypsin, 37°C) and mechanically triturated with a P1000 tip in DMEM (Invitrogen) supplemented with 10% SVF, 0,5mM Glutamax (Invitrogen), Penicillin-Streptomycin 5U/ml (Invitrogen). Cells were plated on 24-well plates (7.10E4 cells per well) on glass coverslips coated with poly-dl-ornithine (Sigma), in Neurobasal® medium (Invitrogen) supplemented with 0,5 mM Glutamax (Invitrogen) and B27 1X (Invitrogen). Experiments were carried out at 15-21 days in vitro (DIV).

#### d-STORM

##### Immunolabelling

Cells were fixed by incubation for 20 min at room temperature in 4% paraformaldehyde in phosphate-buffered saline (PBS) and were rinsed with PBS. After permeabilization in 0,5% Tween 3 % BSA PBS1X, neurons were incubated with the anti-BIN1 99D primary antibody diluted to 1/250 in the permeabilization buffer, for 1h30 at 37°C. The cells were then rinsed and incubated with goat anti-mouse 647 (1/1000, Life Technologies) in 1% BSA PBS1X for 45 minutes at 37°C. At the end, actin was labelled using Phalloidin Alexa 488 (Thermo Fisher), and cells were immerged in the imaging buffer (see below).

##### Optical set-up

Super-localization images were acquired using a Nikon Eclipse Ti inverted microscope combined with a Perfect Focus System and configured for these studies in TIRF excitation. Samples were excited sequentially with 637-nm and 488-nm optically pumped semiconductor lasers (respectively Obis 637 LX 140 mW, Coherent and Genesis MX-STM 500 mW, Coherent). A set of full-multiband laser filters, optimized for 405-, 488-, 561- and 635-nm laser sources (LF405/488/561/635-A-000, Semrock) was used to excite Alexa Fluor 488 and Alexa Fluor 647, as well as for the collection of the resultant fluorescence via a Nikon APO TIRF 60x, NA 1.49 oil immersion objective lens. All images were recorded onto a 512×512-pixels EMCCD camera (iXon 897, Andor), split into two regions of 256×256-pixels area and positioned on the focal plane of an optical telescope for an optimal sampling of the PSF (2.7x magnification, optical pixel size of ∼100 nm). The laser power at 488 nm and 637 nm was 140 mW measured in the BFP. For each frame (∼1500 recorded), the integration time and the EMCCD gain were respectively set to 50 ms and 150.

##### Imaging buffer

During the dSTORM acquisitions, the samples were immerged in an imaging buffer that allows for the fluorophores blinking while reducing their photobleaching (van de Linde et al., 2011; Vaughan et al., 2012). This buffer contains 50-100 mM of ß-mercaptoethylamine (MEA, Sigma-Aldrich) and an oxygen scavenger system (0.5 mg.ml-1 glucose oxidase (Sigma-Aldrich), 40 µg.ml-1 catalase (Sigma-Aldrich) and 10% (w/v) glucose) dissolved in a buffer composed of 100 mM Tris-HCl (Sigma-Aldrich), 1 mM ascorbic acid (Sigma-Aldrich) and 1 mM methyl viologen (Sigma-Aldrich). The pH of the final solution was adjusted to 7.5.

##### dSTORM imaging

To induce the majority of the fluorophores into the dark state, we excited the samples using the laser in an oblique configuration (first 637 nm for Bin1 immunolabeled with Alexa Fluor 647, then 488 nm for F-Actin labeled with phalloidin Alexa Fluor 488). Once the density of fluorescent dye molecules was sufficient (typically, < 1 molecule.µm-2), we switched to a TIRF excitation with an irradiance of 2 kW.cm-2 and activated our home-made real-time super-localization software. This software was written in Python, using PyQt, Numpy and Scipy libraries, and is based on image wavelet-segmentation and centroid determination (Izeddin et al., 2012). This algorithm provides dSTORM images with a localization precision of 15 nm. Drift correction is implemented using the redundant cross-correlation algorithm developed by Wang et al. (Wang et al., 2014). Moreover to improve the viewing quality, each single-molecule detection is added to the final dSTORM image as a 2D Gaussian function with a 15-nm width.

##### Data Analysis

To analyse the location (pre- or post-synaptic) of protein Bin1, we first manually choose synapses of interest using F-actin image. Then, on each synapse Region Of Interest (ROI), we automatically determine contour of dendritic spine (post-synaptic region) thanks to a homemade analysis software written in Python. This contour of dendritic spine can then be used to super-localize pre-synaptic region with a synaptic cleft of 30 nm. Finally, super-localization data of post- and pre-synaptic regions are used to determine the location of Bin1 protein.

#### Tagged Bin1 knockin mouse line establishment

The Bin1-HHStag & Bin1-DMtag mutant mouse lines were established at the Institut Clinique de la Souris (PHENOMIN – ICS, France; www.phenomin.fr/). Bin1-HHStag has a twin tag inserted in exon 19 (C terminal) to tag all isoforms of BIN1. In contrast, Bin1-DMtag has a twin tag inserted in exon 15 to tag only the neuronal isoform of BIN1.

The targeting vector was constructed by Gene Bridges GmbH.

The linearized construct was electroporated in C57BL/6N mouse embryonic stem (ES) cells. After selection, targeted clones were identified by PCR using external primers and further confirmed by Southern blot with 5’ and 3’ external probes.

One positive ES clone was injected into BALB/cN blastocysts, and male derived chimaeras were bred with a FlpO deleter line (Birling et al., 2012) to give germline transmission and remove the selection cassette used for ES cells screening.

#### IP of brain tissues from Tagged BIN1 knockin mice

##### Sample preparation and digestion

Purification and elutions have been done according the protocol of Strep-Tactin©XT:Twin-Strep-tag ©kit, IBA GmbH, using 4 dissected cortical regions of K5288, K5289 and WT 3 months-old C57B6 mice. Immunoprecipitates were prepared as described. After the final elution in desthiobiotin-containing buffer, the proteins were first reduced with 50mM Bond Breaker TCEP solution [(tris(2-carboxyethyl)phosphine), Thermo Fisher Scientific] at RT for 15min and proceeded for digestion using Filter-assisted sample preparation (FASP) method, performed essentially as described (Lipecka et al., 2016; Wiśniewski et al., 2009). Briefly, protein extracts were applied to 30kDa MWCO centrifugal filter units (Microcon, Millipore), mixed with UA buffer (8M urea, 100mM Tris-HCl pH 8.9) and centrifuged. Alkylation was carried out by incubation for 20min in the dark with UA buffer containing 50mM iodoacetamide. Filters were then washed twice with UA buffer followed by two washes with ABC buffer (50 mM ammonium bicarbonate). Finally, trypsin (Promega, France) was added in 1:50 ratio and digestion was achieved by overnight incubation at 37°C. Peptides were collected, vacuum dried and resuspended in 10% acetonitrile, 0.1% formic acid, for LC-MS/MS.

##### Mass spectrometry

For each run, 1 µL were injected in a nanoRSLC-Q Exactive PLUS (Dionex RSLC Ultimate 3000, Thermo Scientific, Waltham, MA, USA). Extracted peptides were resuspended in 0.1% (v/v) trifluoroacetic acid, 10% acetonitrile, and were loaded onto a µ-precolumn (Acclaim PepMap 100 C18, cartridge, 300 µm i.d.×5 mm, 5 µm, Dionex), followed by separation on the analytical 50 cm nano column (0.075 mm ID, Acclaim PepMap 100, C18, 2 µm, Dionex). Chromatography solvents were (A) 0.1% formic acid in water, and (B) 80% acetonitrile, 0.08% formic acid. Peptides were eluted from the column using a gradient from 5% to 40% B over 38 min and were analyzed by data dependent MS/MS, using top-10 acquisition method. Briefly, the instrument settings were as follows: resolution was set to 70,000 for MS scans and 17,500 for the data dependent MS/MS scans in order to increase speed. The MS AGC target was set to 3.10^6^ counts with a maximum injection time of 200 ms, while MS/MS AGC target was set to 1.10^5^ with a maximum injection time of 120 ms. Dynamic exclusion was set to 30 sec. Each sample was analyzed in three to five biological replicates.

##### Data Processing Following LC-MS/MS acquisition

Raw MS files were processed with the MaxQuant software version 1.5.3.30 and searched with Andromeda search engine against the Mus musculus Uniprot KB/Swiss-Prot v.06/2016. To search parent mass and fragment ions, we set an initial mass deviation of 4.5 ppm and 20 ppm respectively. The minimum peptide length was set to 7 aminoacids and strict specificity for trypsin cleavage was required, allowing up to two missed cleavage sites. Carbamidomethylation (Cys) was set as fixed modification, whereas oxidation (Met) and N-term acetylation were set as variable modifications. The false discovery rates (FDRs) at the protein and peptide level were set to 1%. Scores were calculated in MaxQuant as described previously (Cox and Mann, 2008). The reverse and common contaminants hits were removed from MaxQuant output. Proteins were quantified according to the MaxQuant label-free algorithm using LFQ intensities (Cox and Mann, 2008; Luber et al., 2010). Protein quantification was obtained using at least 2 peptides per protein.

Five biological replicates were analysed Bin1, Four for Bin1D and three WT (used as control). Statistical analysis between Bin1 and all the others was performed with Perseus software (version 1.5.5.3) freely available at www.perseus-framework.org. For statistical comparison, the LFQ (Label-free Quantification) data were transformed in log2, the matrix was filtered to keep proteins identifies at least four times in at least the Bin1 or Bin1D group. The remaining missing data were imputed by creating a Gaussian distribution of random numbers with a standard deviation of 30% relative to the standard deviation of the measured values and 1.8 standard deviation downshift of the mean to simulate the distribution of low signal values. We analysed the data by t-test using FDR correction, and represented the data on volcano plot (T test FDR<0.01; S0=0.2).

#### Analysis of IP data using WebGeslalt

Data from immunoprecipitation analysis were analyzed with WebGestalt (WEB-based Gene SeT AnaLysis Toolkit) that is a functional enrichment analysis web tool (Wang et al., 2017).

#### Synaptosomal preparation

Fresh cortices and hippocampus from 2-5 months-old OF1 male mice were dissected and homogenized in 30 ml Buffer H 1X (0.32 M Sucrose, 5mM Hepes, 1mM EDTA) using a glass douncer (9 strokes, 800 rpm). After centrifugation (7 min, 2600 rpm, 4°C), the supernatant, containing cytosolic fraction and synaptosomes, was centrifuged again (20 min, 5000 rpm,4°C) resulting in a pellet containing the synaptic vesicles. The latter was resuspended in Buffer H 1X, put on a 5%-10%-23% Percoll gradient (2m of suspension per gradient), and centrifuged (11 min, 18 000 rpm, 4°C). The synaptosomal fraction, between the 10% and the 23% phases, was centrifuged (20 min, 4000 rpm, 4°C) and the resulting pellet was resuspended in lysis buffer (50 mM NaCl, 50 mM Tris pH8, 1% NP40, 0.5% Na Deoxycholate, Phosphatase and Protease inhibitors).

#### Coimmunoprecipitation assays

Protein samples from synaptosomal protein fraction (0.4 mg) were precleared with 30µl of magnetic beads (protein G coated Dynabeads, Novex) during 45 min under rotation at 4°C. 5µg of Primary antibodies anti-BIN1 99D or Mouse IgG were bound to 50 µl protein G-coupled beads during 45 minutes at 4°C. Precleared lysates were incubated with the antibody-bead complexes overnight under rotation at 4°C. Supernatants were collected and after three washes (150 mM NaCl 1% NP40 10mM Tris pH 7,5 buffer), beads were eluted with 2X SDS polyacrylamide gel electrophoresis sample buffer 10 min at 95°C and analysed with western blotting.

#### LC/MS/MS from synaptosomes

##### Samples preparation

LC/MS/MS experiments were carried out on BIN1 immunoprecipitation from synaptosomal fraction. For each experiment, 10 mg of proteins were used. For the first experiment the whole amount of proteins was processed in one time. For the second and third experiments, it was divided in 25 experiments and eluates were pooled at the end. Beads G were cross-linked to BIN1 antibody or IgG Mouse. Precleared lysates were incubated with the antibody-bead complexes overnight (first and second experiment) or 2h (third experiment) under rotation at 4°C. Supernatants were collected and after three washes (150 mM NaCl 1% NP40 10mM Tris pH 7,5 buffer) and beads were eluted with PBS1X SDS 0,5 % (first and second experiment) or PBS1X SDS 3% (third experiment) 10 min at 95°C.

LC/MS/MS was carried out by Dualsystems Biotech AG, Schlieren, Switzerland (www.dualsystems.com). Samples for mass spectrometry analysis have been processed according a filter-aided sample preparation (FASP) protocol (Wiśniewski et al., 2009). In short, eluates from affinity purifications have been denatured with 6M Urea, reduced with 50 mM DTT for 30 minutes at 37°C and subsequently alkylated with 200 mM iodoacetamide for 30 minutes at room temperature. The samples have then been loaded onto a standard filtration device (30 kDa MWCO; VivaCon 500, Sartorius) and centrifuged at 14’000 x g for 15 minutes at 20°C. To exchange the buffer, 8M Urea was added to the samples and the centrifugation step was repeated for a total of four times. Ammonium bicarbonate together with 1 µg of Trypsin (Proteomics grade, Sigma) was added to the samples for an overnight protein digestion at 37°C. Tryptic peptides have been collected by centrifugation with three subsequent 40 mM ammonium bicarbonate washes. The tryptic peptides have been acidified to 1% trifluoroacetic acid (Sigma) and desalted by MicroSpin C18 Silica columns (The Nest Group. Inc.) according the manufacturers protocol.

##### Liquid chromatography tandem mass spectrometry (LC-MS/MS) analysis)

Tryptic peptides have been analyzed on a LTQ Orbitrap XL (Thermo Scientific) mass spectrometer fitted with a nanoelectrospray ion source (Thermo Scientific). The peptide samples were separated by reverse phase chromatography on a high performance liquid chromatography column with 75 µm inner diameter (Michrom Bioresources), packed with a 10 cm stationary phase (Magic C18AQ, 200 Å, 3 m, Michrom Bioresources) connected to an EASY-nLC 1000 (Thermo Sientific) nano flow HPLC. Peptides were loaded onto the column in 95% buffer A (98% H_2_O, 2% acetonitrile, 0.1% formic acid) and eluted with 300 nl/min over a 60 minute linear gradient from 5-35% buffer B (2% H_2_O, 98% acetonitrile, 0.1% formic acid). Mass spectra were acquired in a data dependent mode. High resolution MS scans were acquired in the Orbitrap (60’000 FWHM, target value 10^6^) to monitor peptide ions in the mass range of 350-1650 *m/z*, followed by collision induced dissociation MS/MS scans in the ion trap (minimum signal threshold 150, target value 10^4^) of the five most intense precursor ions.

##### MS data analysis and quantification

The raw data was analyzed with MaxQuant (version 1.3.0.5) as described (Cox and Mann, 2008) using the Andromeda search engine against UniProt protein database of Mus musculus, concatenated with the sequences of common contaminants. A reverse decoy database approach was used to filter the protein identifications to a false discovery rate of ≤1%.

#### Proximity ligation assays (PLA) and microscopy

We followed the protocol described in (Söderberg et al., 2006). Cells were fixed by incubation for 20 min at room temperature in 4% paraformaldehyde in phosphate-buffered saline (PBS) and PLA was realized according to the instructions of the manufacturer (DuoLink, Sigma). Primary antibodies used were all incubated in 0,5% Tween, 3% BSA PBS1X for 1h30 at 37°C (Dilutions described in Supplementary Table). At the end of the PLA protocol, actin was labelled using Phalloidin Alexa 488 (Thermo Fisher). For the analysis of PLA interactions points, cells were scanned using the laser scanning confocal microscope (Leica, SP5 from PICPEN imagery platform Centre de Psychiatrie et Neuroscience) at 63× magnification, zoom 3.43 to get optimal pixel size, and 0.25 µm z-stacks. PLA interaction number was counted using volocity software on 3D-reconstructions on 70 μm long dendritic segments and normalized by actin volume. One 70 μm dendritic segment per neuron was analyzed.

When PLA was associated with PSD-95 or Gephyrin immunofluorescence, a step of pre-permeabilization of 30 minutes in PBS1X Tween 0,1% was performed before the PLA protocol. Immunoflurescence was performed directly after the last wash of the PLA protocol. Primary and secondary antibodies used are described in Supplementary Table. Cells were scanned as described for the PLA assay. Pearson coefficient was performed using the Volocity software on one dendritic segment per neuron.

#### RT-qPCR

Reverse transcription (RT) was carried out using the Maxima First Strand Kit (Fermentas). The cDNAs generated were amplified by real-time PCR, using probes (obtained from Eurogentec) labeled at their 5′ ends with a fluorogenic reporter dye and at their 3′ ends with a quencher dye (TAMRA). Sequences of primers and probes are available on request. PCR assays had a final reaction volume of 20 µl, and contained 2 U of *Taq* polymerase (Master Mix, Applied Biosystems), 10 µM primers and fluorogenic probe. PCR was carried out over 40 cycles of 95°C for 15 s, 60°C for 1 min, 50°C for 1 min. We used the Opticon2 sequence detection system (MJ Research/Biorad), with the Opticon Monitor software for data analysis. For each group, the cDNAs synthesized from total RNA were serially diluted to cover the 0.008–50 ng range for specific mRNAs and 0.008–2.5 ng range for 18S rRNAs. These serial dilutions were used to construct standard curves for 18S and for each gene of interest and to calculate the amounts of RNA for 18S and the genes of interest corresponding to the PCR products generated from individual cDNAs for each experimental group. Each Q-PCR signal was normalized with respect to 18S.

#### Active Rac1 Detection in hippocampus

Protein samples were prepared from microdissection of 10 hippocampi of juvenile mice incubated in RIPA lysis buffer (Santa Cruz sc-24948). Active Rac1 Detection was performed using Active Rac1 Detection Kit of Cell Signalling Technology. Briefly, a GTP-bound GTPase pull-down was performed and the eluted samples revealing Rac1 activation were analysed by western blot using a Rac1 mouse antibody provided by the kit.

#### Aβ ELISA

The Aβ extracted was quantified with the MSD V-PLEX Aβ Peptide Panel 1 (4G8) kit (Meso Scale Diagnostics, Rockville, USA). Kit provides assay-specific components for the quantitative determination of Aβ38, Aβ40, and Aβ42 in human and rodent samples. ELISA was performed according to the kit manufacturer’s instructions in each case. We used supernatant of human SH-SY5Y neuroblastoma cells that overexpress APP with the Swedish mutation as positive control.

