## Supplemental Figures for "BIN1 genetic risk factor for Alzheimer is sufficient to induce early structural tract alterations in entorhinal-hippocampal area and memory-related hippocampal multi-scale impairments"

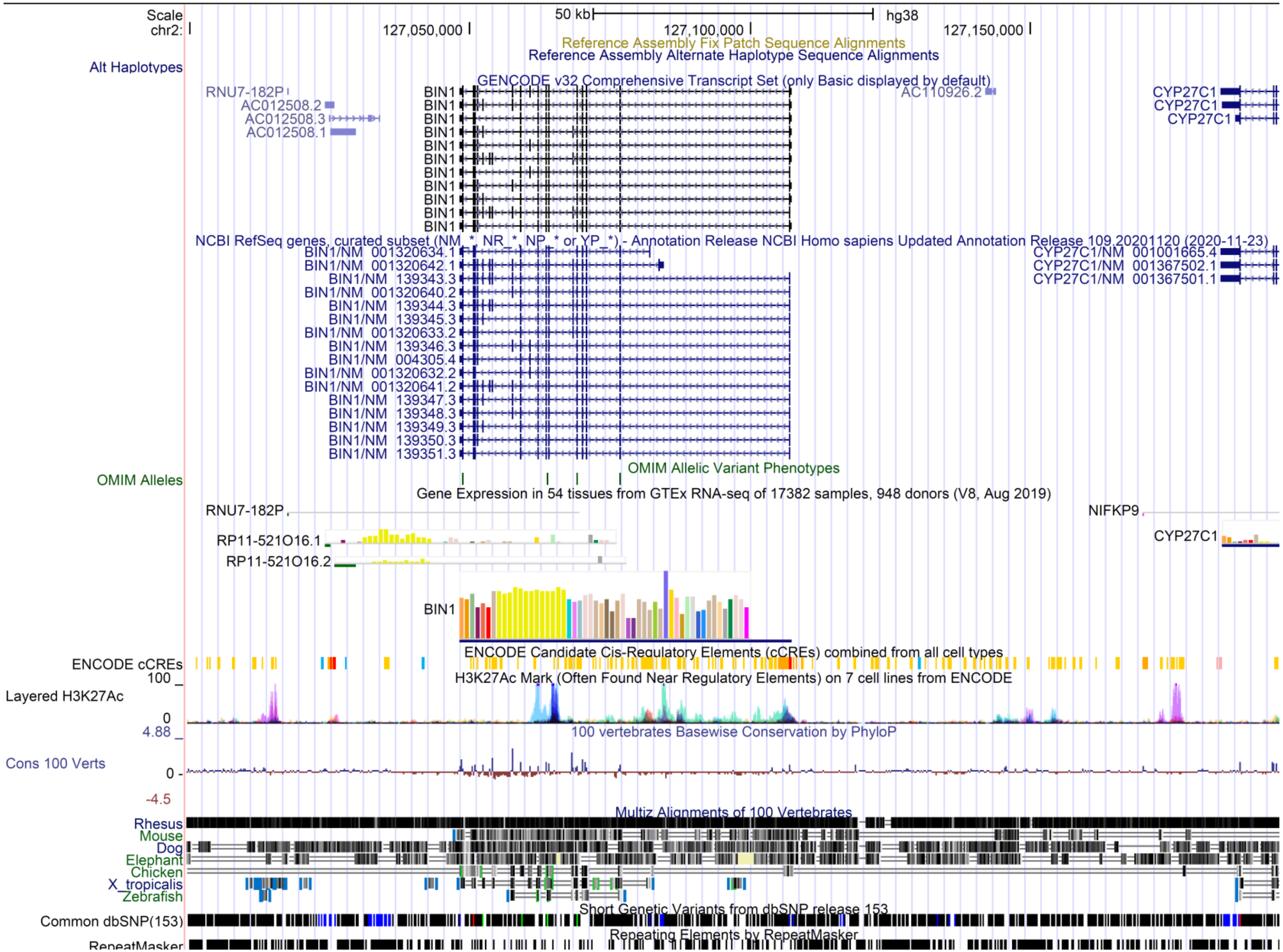
Scale
chr2:
50 kb
hg38
127,050,000
127,100,000
127,150,000
Reference Assembly Fix Patch Sequence Alignments
Reference Assembly Alternate Haplotype Sequence Alignments
Alt Haplotypes
GENCODE v32 Comprehensive Transcript Set (only Basic displayed by default)
RNU7-182P
AC012508.2
AC012508.3
AC012508.1
BIN1
AC110926.2
CYP27C1
NCBI RefSeq genes, curated subset (NM_*, NR_*, NP_* or YP_*) - Annotation Release NCBI Homo sapiens Updated Annotation Release 109.20201120 (2020-11-23)
BIN1/NM_001320634.1
BIN1/NM_001320642.1
BIN1/NM_139343.3
BIN1/NM_001320640.2
BIN1/NM_139344.3
BIN1/NM_139345.3
BIN1/NM_001320633.2
BIN1/NM_139346.3
BIN1/NM_004305.4
BIN1/NM_001320632.2
BIN1/NM_001320641.2
BIN1/NM_139347.3
BIN1/NM_139348.3
BIN1/NM_139349.3
BIN1/NM_139350.3
BIN1/NM_139351.3
CYP27C1/NM_001001665.4
CYP27C1/NM_001367502.1
CYP27C1/NM_001367501.1
OMIM Allelic Variant Phenotypes
OMIM Alleles
Gene Expression in 54 tissues from GTEx RNA-seq of 17382 samples, 948 donors (V8, Aug 2019)
RNU7-182P
NIFKP9
RP11-521O16.1
RP11-521O16.2
CYP27C1
BIN1
ENCODE Candidate Cis-Regulatory Elements (cCREs) combined from all cell types
ENCODE cCREs
100
H3K27Ac Mark (Often Found Near Regulatory Elements) on 7 cell lines from ENCODE
Layered H3K27Ac
0
4.88
100 vertebrates Basewise Conservation by PhyloP
Cons 100 Verts
0
-4.5
Multiz Alignments of 100 Vertebrates
Rhesus
Mouse
Dog
Elephant
Chicken
X_tropicalis
Zebrafish
Short Genetic Variants from dbSNP release 153
Common dbSNP(153)
Repeating Elements by RepeatMasker
RepeatMasker

Supplementary Fig.1. **Position of human BAC RP11-437K23 in the human genome (version GRCh38)**

BAC RP11-437K23 spans over 195 kb. This BAC contains only the complete protein coding gene BIN1. All coding exons were PCR amplified and confirmed by Sanger sequencing.

The putative 5’ and 3’ ends of the RP11-437K23 are located at Chr2: 127,194,398 and 126,999,436 bp (GRCh38) respectively.

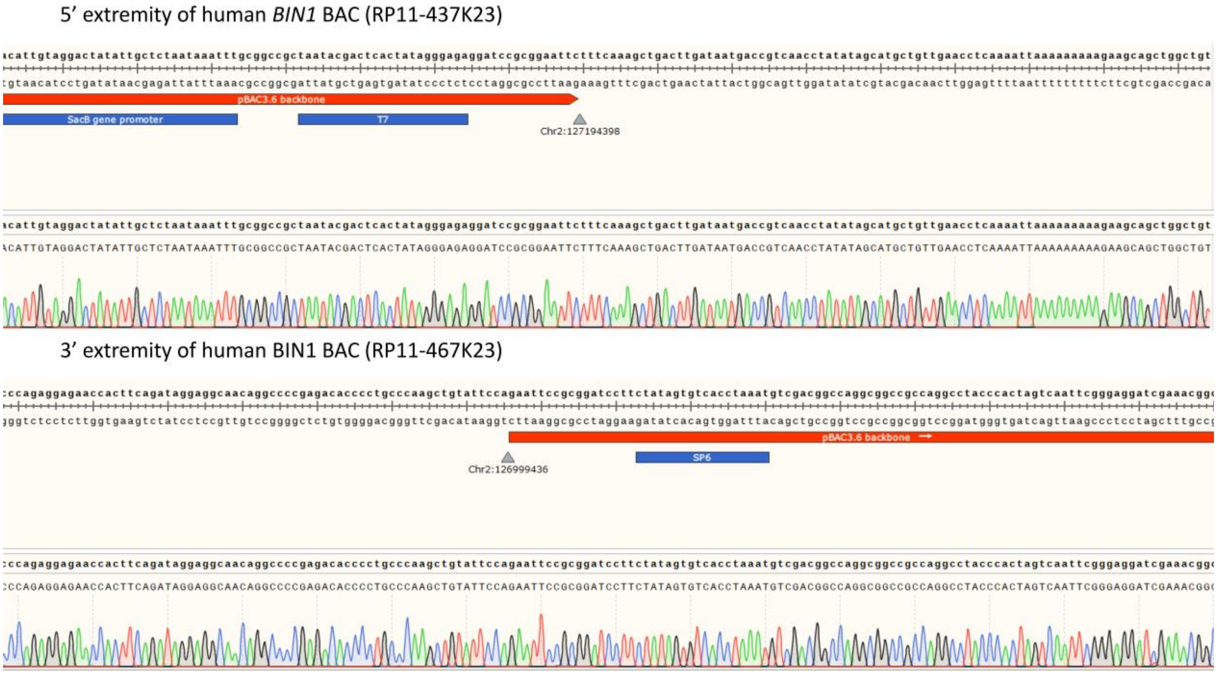


Supplementary Fig.2. **Analysis of *hBIN1* BAC ends.**

The extremities of BAC RP11-437K23 were sequence after PCR amplification from the extremity of the backbone (pBACe3.6) to the supposed extremity of the human sequence. The Sanger sequencing results of both the 5' and 3' extremity of the BAC are show in this figure.

The human sequence spans from Chr2:126,999,436-127,194,398 (alignement against GRCh38.p13 Ensembl release 99). The BAC RP11-437K23 was annotated on the former human genome alignment (GRCh37 Full Feb 2014 archive with BALAST, VEP and BioMart) at the position Chromosome 2: 127,761,089-127,941,604 (https://grch37.ensembl.org/Homo_sapiens/Location/View?db=core;g=ENSG00000136717;r=2:127761089-127941604)

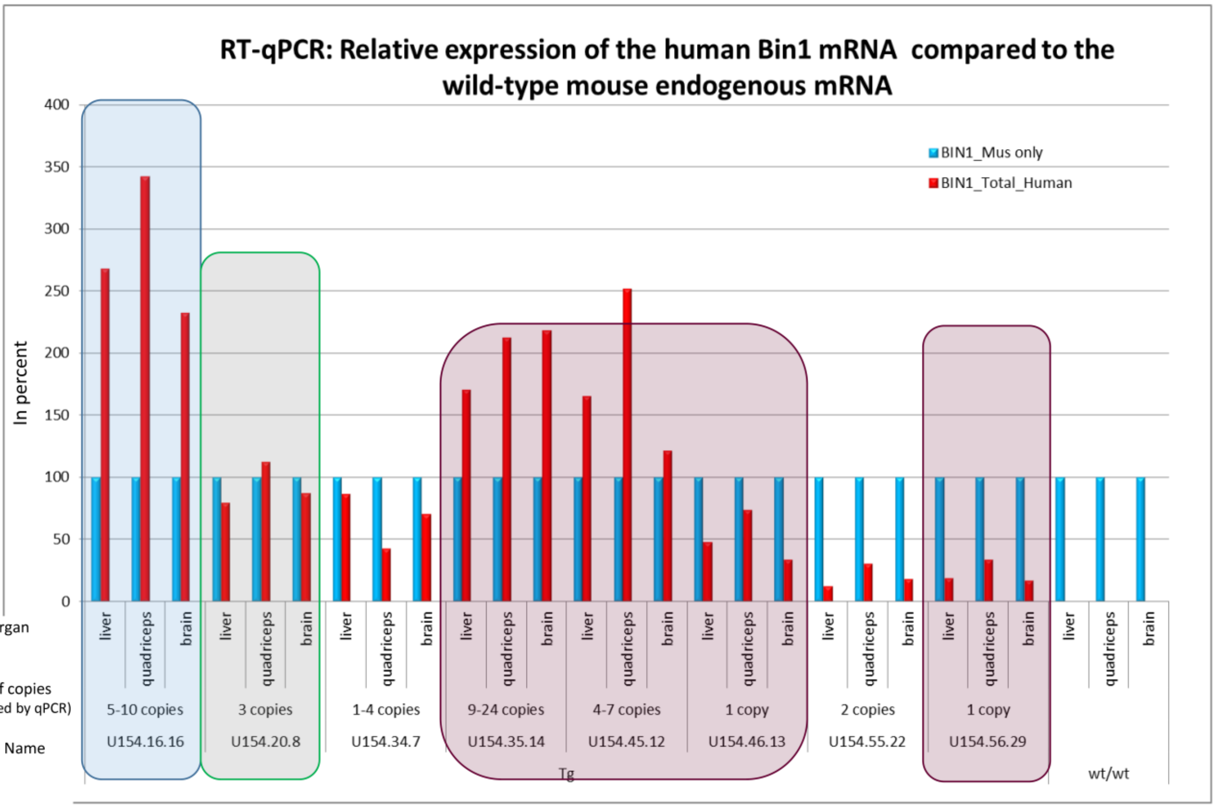


Supplementary-Fig.3

Supplementary Fig.3. **Relative expression of human BIN1 mRNA compared to the expression of endogenous mouse Bin1 mRNA as analyzed by RT-qPCR, for the 8 *hBIN1* lines**

Supplementary Fig.3. **Relative expression of human BIN1 mRNA compared to the expression of endogenous mouse Bin1 mRNA as analyzed by RT-qPCR, for the 8 *hBIN1* lines**

Relative expression of human BIN1 mRNA for the different hBIN1 line is compared to the expression of endogenous mouse Bin1 mRNA as analyzed by RT-qPCR. Number of BAC integrated is indicated

### *3 months MD ($m^2/s$)*

*Global ROIs (whole brain) :* *Hippocampal-Entorhinal area :*

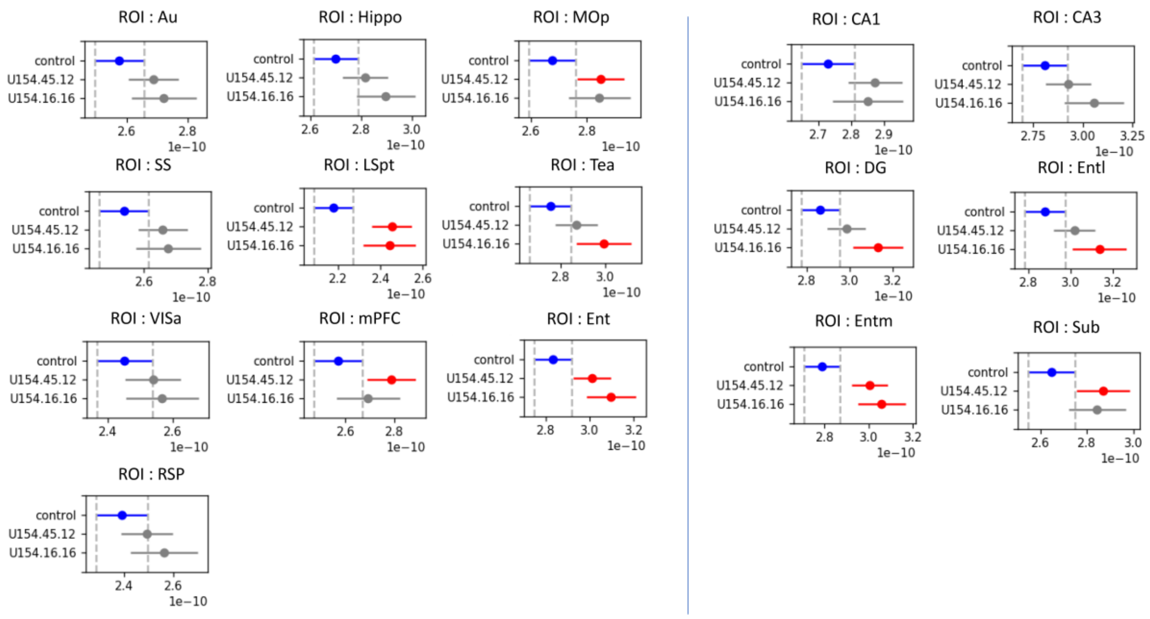


### *6 months MD ($m^2/s$)*

*Global ROIs (whole brain) :* *Hippocampal-Entorhinal area :*

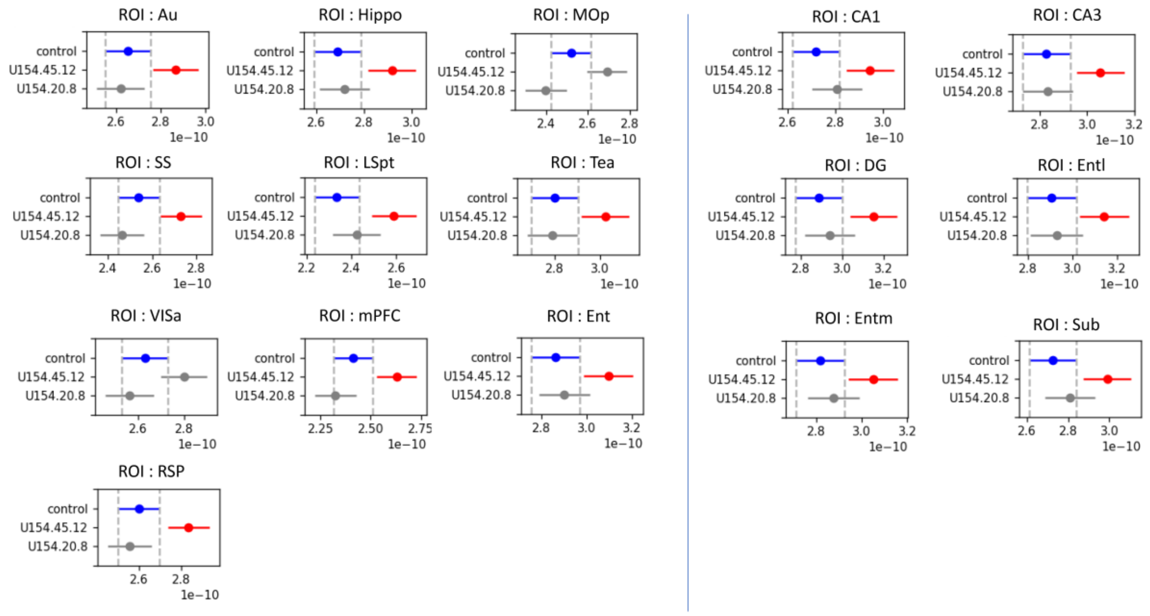


Supplementary Fig.4. **Comparison of three distinct *hBIN1* lines: Commonality of MD scalar changes for line 2 and line 3 and absence of changes for line 1.**

Supplementary Fig.4. **Comparison of three distinct *hBIN1* lines: Commonality of MD scalar changes for line 2 and line 3 and absence of changes for line 1.**

Confidence intervals of the MD for each ROI obtained in each group with Tukey HSD test at 3 months. The three groups at 3 months are WT (control), hBIN1 line 3 (U154.16.16), hBIN1 line 2 (U154.45.12). The significance level was set to alpha = 0.05, FWER-corrected. The blue lines are the values of the reference group, here the WT group. The MD values of each hBIN1 line are compared to the WT group. Significant differences are colored in red. The two hBIN1 lines show the same tendency at 3 months (MD increase). The MD values of each hBIN1 line are compared to the WT group. hBIN1 line 2 has increased MD in the whole brain with all hippocampal-entorhinal areas affected.

Confidence intervals of the MD for each ROI obtained in each group with Tukey HSD test at 6 months. The three groups at 6 months are WT (control), hBIN1 line 2 (U154.45.12), hBIN1 line 1 (U154.20.8). The significance level was set to alpha = 0.05, FWER-corrected. The blue lines are the values of the reference group, here the WT group. The MD values of each hBIN1 line are compared to the WT group. hBIN1 line 2 has increased MD in the whole brain with all hippocampal-entorhinal areas affected, while hBIN1 line 1 shows no significant difference with WT, either in hippocampal-entorhinal area or in global ROIs.

Abbreviations: Au : primary auditory area ; Hippo : hippocampus ; MOp : primary motor area ; SS : somatosensory area ; LSpt : lateral septal nucleus ; Tea : temporal association areas ; VISa : visual areas ; mPFC : medial prefontal cortex ; Ent : entorhinal cortex ; RSP : retrosplenial cortex ; DG : dentate gyrus ; Entl : entorhinal area, lateral part ; Entm : entorhinal area, medial part ; Sub : subiculum

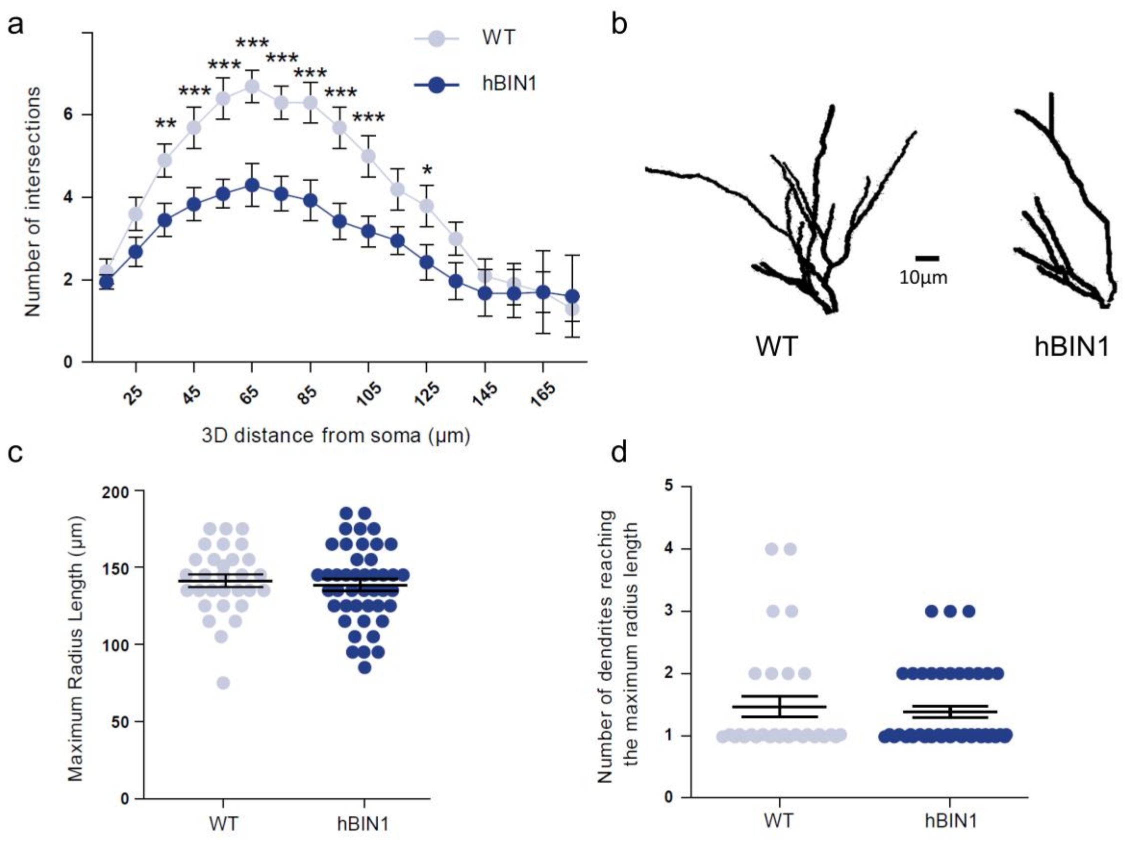


Supplementary Fig.5. **Dendritic tree of granule cells is simplified in hBIN1 mice**

Supplementary Fig.5. **Dendritic tree of granule cells is simplified in hBIN1 mice**

To evaluate dendritic arborisation of granule cells, we performed **Golgi-Cox** staining on fresh brain slices. Labeling was done on 4 animals per genotype at 3 months of age. After confocal imaging, we selected granule cells (n=30 for wt and n= 44 for Tg) for a 3D reconstruction of dendritic tree. (a) Using Scholl analysis, we quantified the number of neuronal branches interecting virtual spheres. hBIN1 had less ramifications compared to WT. (b) Examples of hBIN1 and WT DG granule cells. (c-d) Absence of effect on either maximum radius length (c) and number of dendrites reaching the maximum radius length (d).

*Statistics: Kurtosis and Skewness normality test, ANOVA repeated measures,Bonferroni post hoc test, *** p<0,05; ** p<0,01; * p<0,001).*

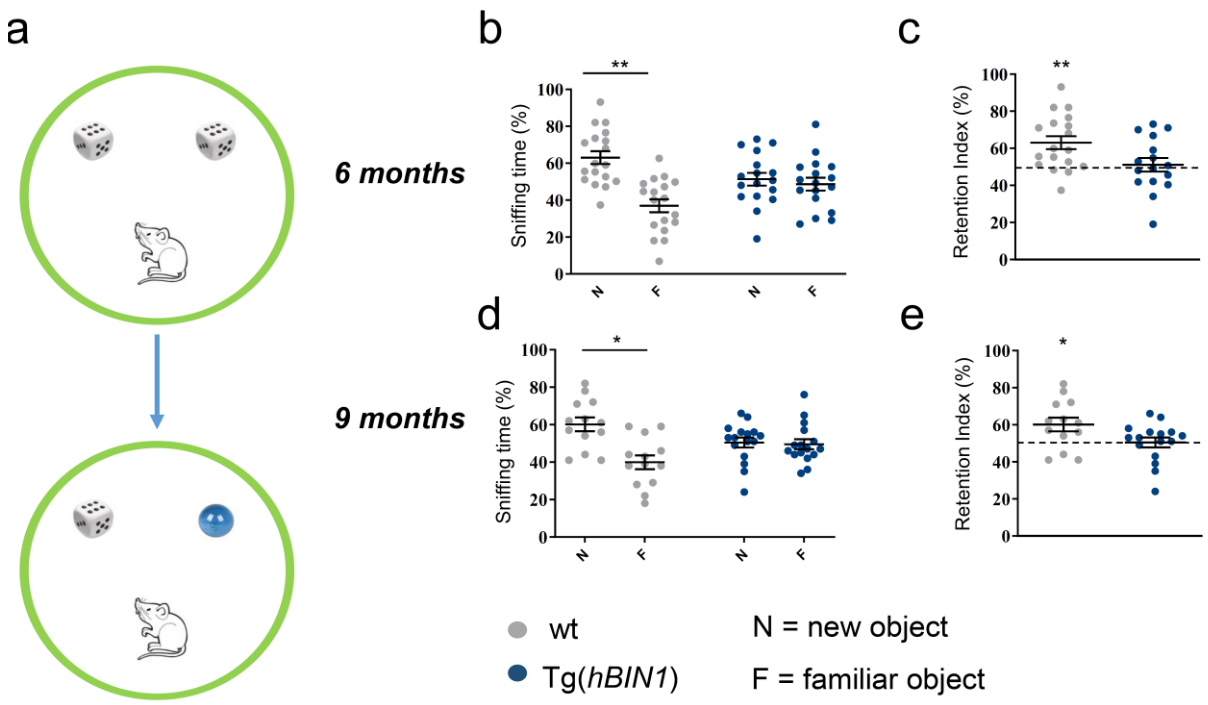


<u>Supplementary Fig. 6.</u> **Object recognition memory is altered in transgenic human Bin1 mice at 6 and 9 months of age.**

Schematic representation of the Novel Object Recognition paradigm (a). Animals were presented with a pair of identical objects (marble or dice) until they had explored the objects more than 3 seconds, in a maximum period of 10 min starting from the first sniffing behavior. In the second trial (test phase, 1h later), one of the familiar objects was changed for a new one, and the animals were left in the arena during 10 min. The sniffing time (reported as percentage of exploration in panel b and d) and the deduced discrimination index (c and de for both wildtype and Tg(hBIN1) staggered at 6 and 9 months of age are shown. Control mice were able to discriminate familiar from novel object, as illustrated by the significative amount of time spent on the new object compare to the familiar one, as well as the discrimination index statistically above the chance (50%), whereas the Tg(hBIN1) could not.

The LOAD candidate gene BIN1 induced short-term memory deficits from 6 months of age.

Data are represented as scattered individual dots plot, with mean + S.E.M; Stastistics: student t-test, new versus familiar object for both genotypes and One sample t-test for the discrimination index versus 50%, are represented by * ($p<0.05$) and ** ($p<0.01$).

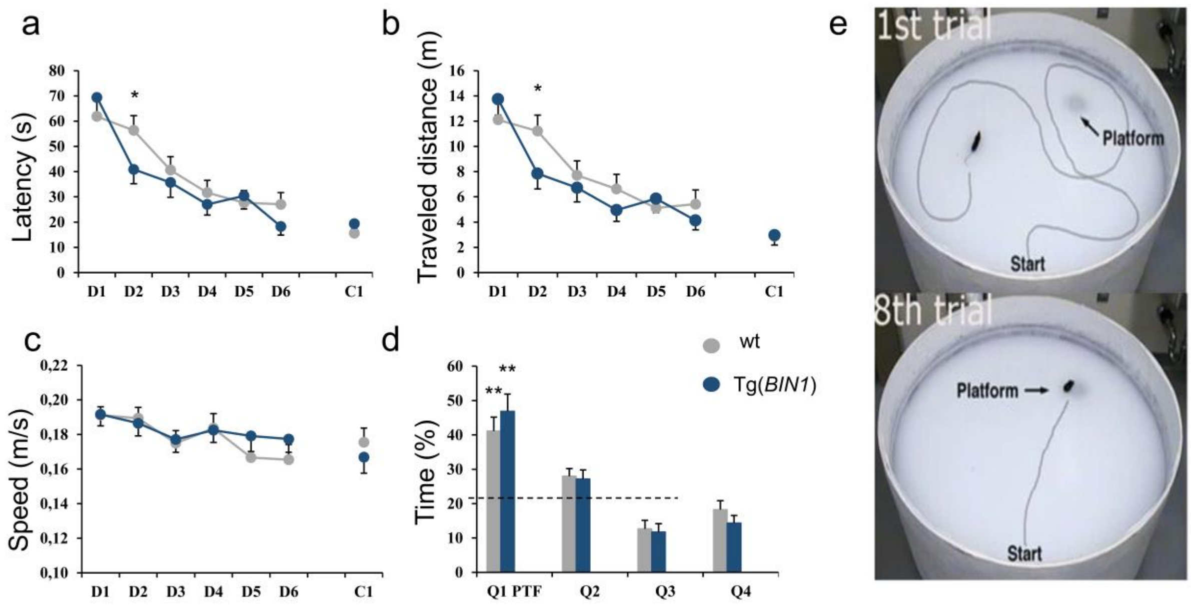


Supplementary Fig. 7. **Analysis of spatial episodic memory tested using Morris Water Maze at month 9.**

Learning of navigation to the hidden platform was not modified between *hBIN1* mice and WT mice as evidenced by three parameters: Latency (**a**), Traveled distance (**b**) and Speed (**c**). D1 to D5 are training days, D6 is the probe trial, D7 is rest and D8 is the day where cued trial is done (C1).

Memory of platform location was not different between Tg(hBIN1) mice and WT mice as evidenced by quadrants occupancy (**d**) during probe trials on day 6. The platform (PTF) is located in quadrant Q1 with Q2, Q3 and Q4 as other quadrants. Both WT and *hBIN1* mice spent more time in the reference quadrant compare to hazard (25%).

Examples of trajectory at first and 8th trials are illustrated in (**e**).

D1 Data are presented as mean + S.E.M; Statistics: Two-way ANOVA repeated measures for the MWM parameters and One sample t-test for the probe trial versus 25%; asterisks denote significant differences (*p < 0.05; **p < 0.01).

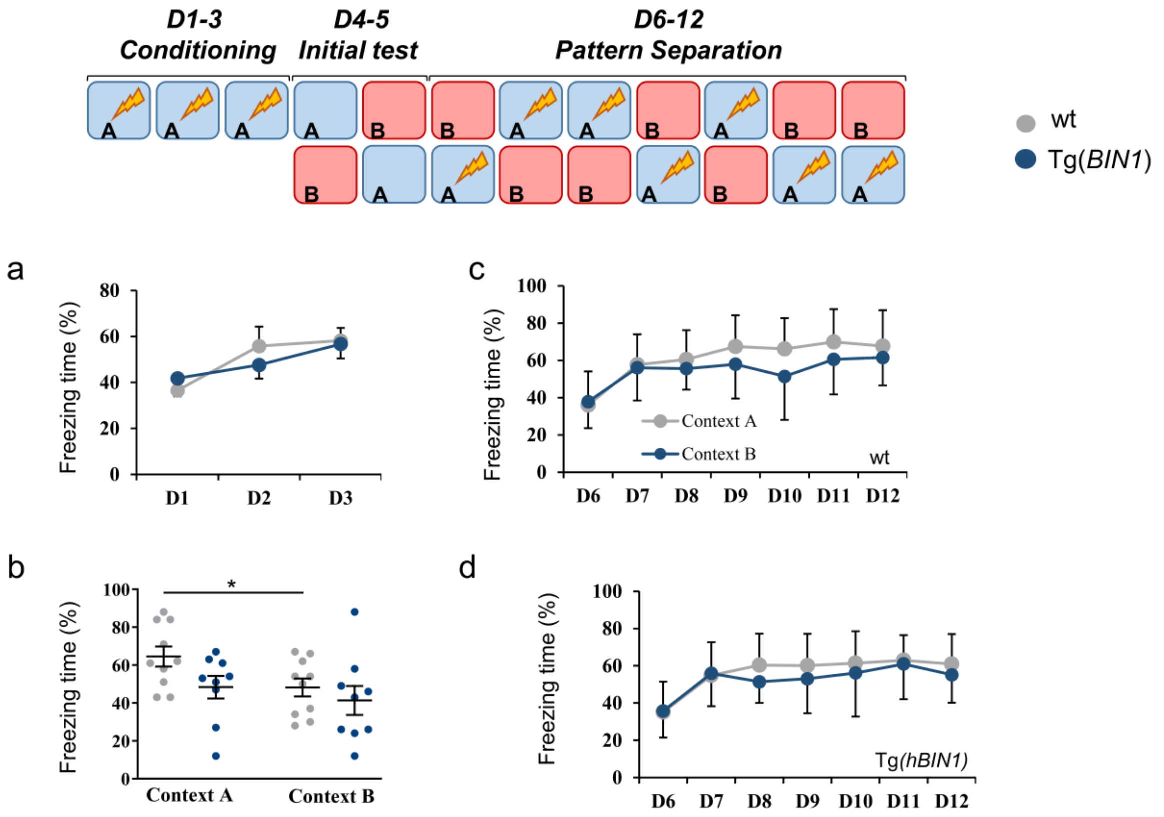


Supplementary Fig. 8. **Tg(hBIN1) displayed impaired contextual memory at 15 months.**

Contextual discrimination protocol is schematized in (a). For the first 3 days of conditioning, mice visited only

chamber A and each day received a single footshock. Freezing was measured once in chamber A and once in chamber B over the subsequent 2 days. During days 6 to 12, mice visited each chamber daily (receiving a shock in one of the two), and freezing was assessed during the first 3 min in each chamber. Both wt and Tg(*hBIN1*) genotypes showed conditioning learning 15 months

(**a**). On D4-5, wild type animals shown a freezing time increase in Context A . Context A corresponds to context where animals received unconditioned stimulus (shock)

**(b**). In contrast, Tg(*hBIN1*) mice are unable to discriminate the two environments A and B (b).

FOR the pattern separation paradigm, both groups behaved the same in response to the repetitive stimuli exposures (**c and d**).

Data are represented as scattered individual dots plot (b), or mean + S.E.M (a, c and d); Statistics: Two-way ANOVA repeated measures for the panel a c and d, and Student t-test, context A versus B for each genotype; differences are represented by * ($p<0.05$).

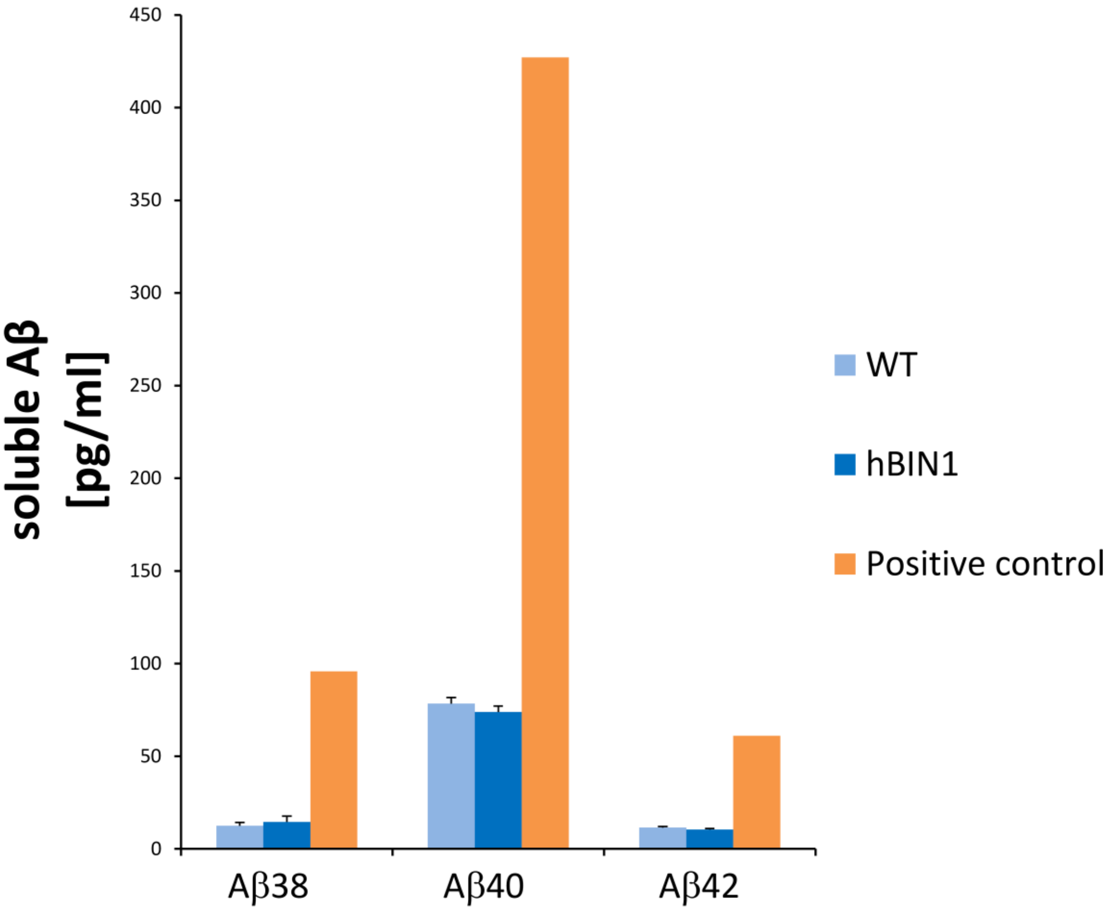


Supplementary Fig.9. **Absence of_Abeta changes in hBIN1 mice.**

Abeta 38, 40 and 42 levels were measured in brain of 12-month-old hBIN1 mice (n=10) and controls (*n* = 8), using multiplex MSD V-PLEX Ab Peptide panel 1 (4G8) immunoassay kit (MesoScaleDiscovery). No changes in these levels have been identified. Positive controls used supernatant analysis of human SH-SY5Y neuroblastoma cells overexpressing human APP with the Swedish mutation. Bars indicate means ± SEM.

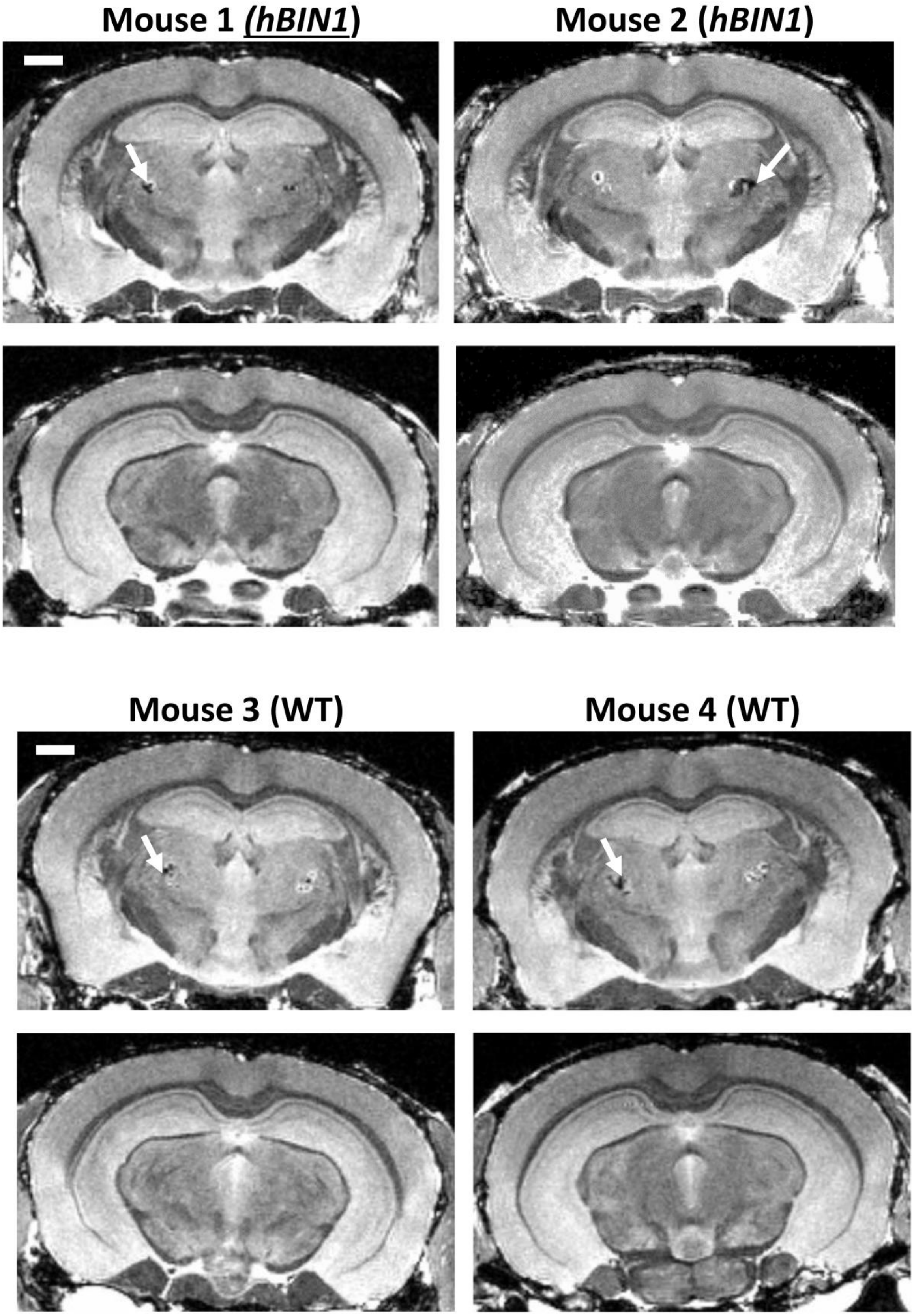


Supplementary Figure 10. **Absence of hypo-densities in neocortex and hippocampus at month 18 in hBIN1 mice.**

Supplementary Figure 10. **Absence of hypo-densities in neocortex and hippocampus at month 18 in hBIN1 mice.**

Comparison between 18 months old BIN1 (left panels) and WT (right panels) mice at different coronal slice levels.

The acquisitions were performed ex vivo (see manuscript for details regarding sample preparation) using 3D T2-weighted TurboRARE acquisition with isotropic resolution of 60 mm and effective echo time of 26 ms.

Similar approaches have been used to detect Abeta plaques as hypo-densities in two genetic mouse models of familial forms of Alzheimer's disease (Zhang et al., Detection of Amyloid Plaques in Mouse Models of Alzheimer's Disease by Magnetic Resonance Imaging. Magnetic Resonance in Medicine 51:452– 457, 2004). Hypointense regions, indicated by white arrows, are detected only in the thalamus in both BIN1 and WT and are due to normal aging.

Scale bar: 1mm.

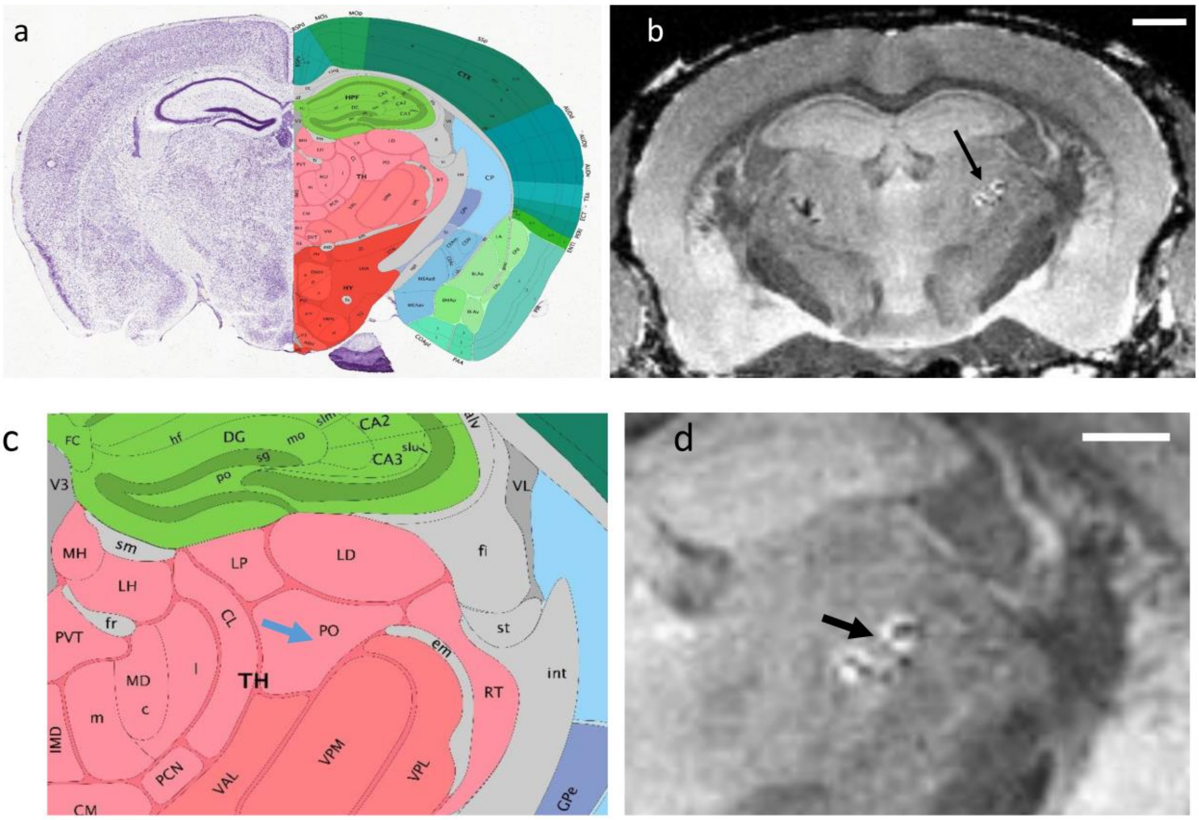


<u>Supplementary Figure 11.</u> **Thalamic hypo-densities in both wild-type and hBIN1 mice at month 18.**

Comparison between Allen Brain reference atlas coronal section (a) and MRI section (b) of a 18 months old wt mouse brain. Hypodensities are similarly located for hBIN1 sections. Enlargement of these sections indicated that hypo-densities are located in PO part of the thalamus and indicated by blue and black arrows in (c) et (d), respectively.

Note that cerebral microbleeds have been reported in thalamus of aged mice brain (Taylor EN, Huang N, Wisco J, Wang Y, Morgan KG, Hamilton JA. The brains of aged mice are characterized by altered tissue diffusion properties and cerebral microbleeds. J Transl Med. 2020 Jul 8;18(1):277.

Scale bar: 1mm for (a) and (b), 300µm for (c) and (d).

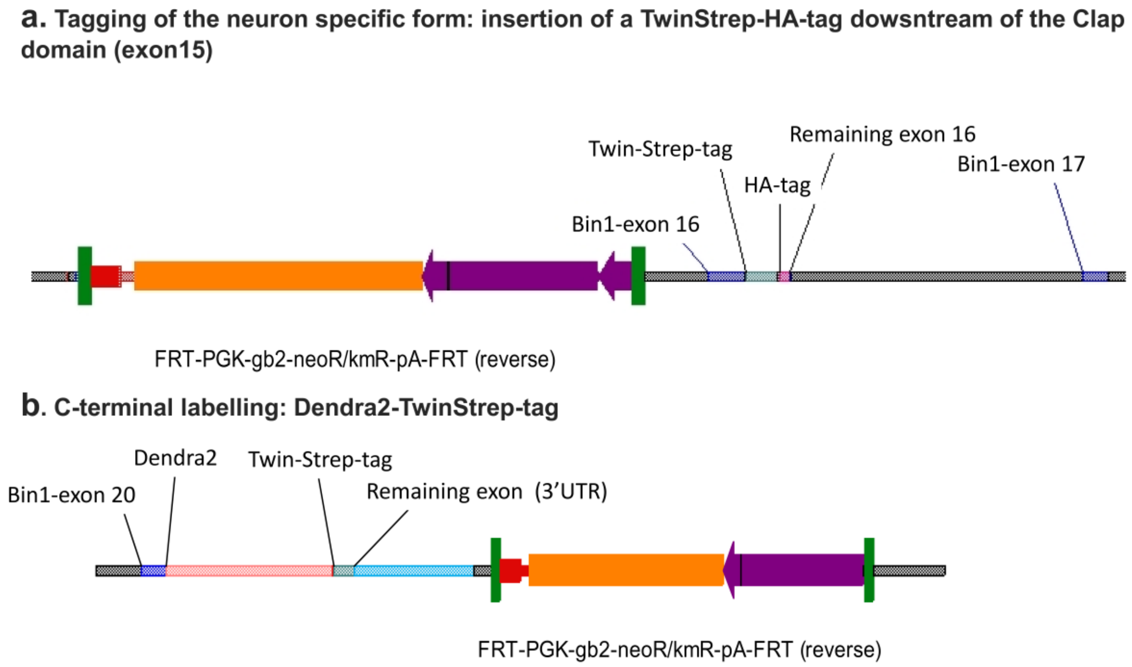


Supplementary Fig. 12 . **Map of the linearized targeting vectors for tagged Bin1 knockin mice.**

Supplementary Fig. 12. .**Map of the linearized targeting vectors for tagged Bin1 knockin mice.**

a: Schematic of Tagging of the neuron specific form: insertion of a TwinStrep-HA-tag dowsntream of the Clap domain (exon16).

b: Schematic of Tagging of all isoforms by C-terminal labelling: Dendra2-TwinStrep-tag.

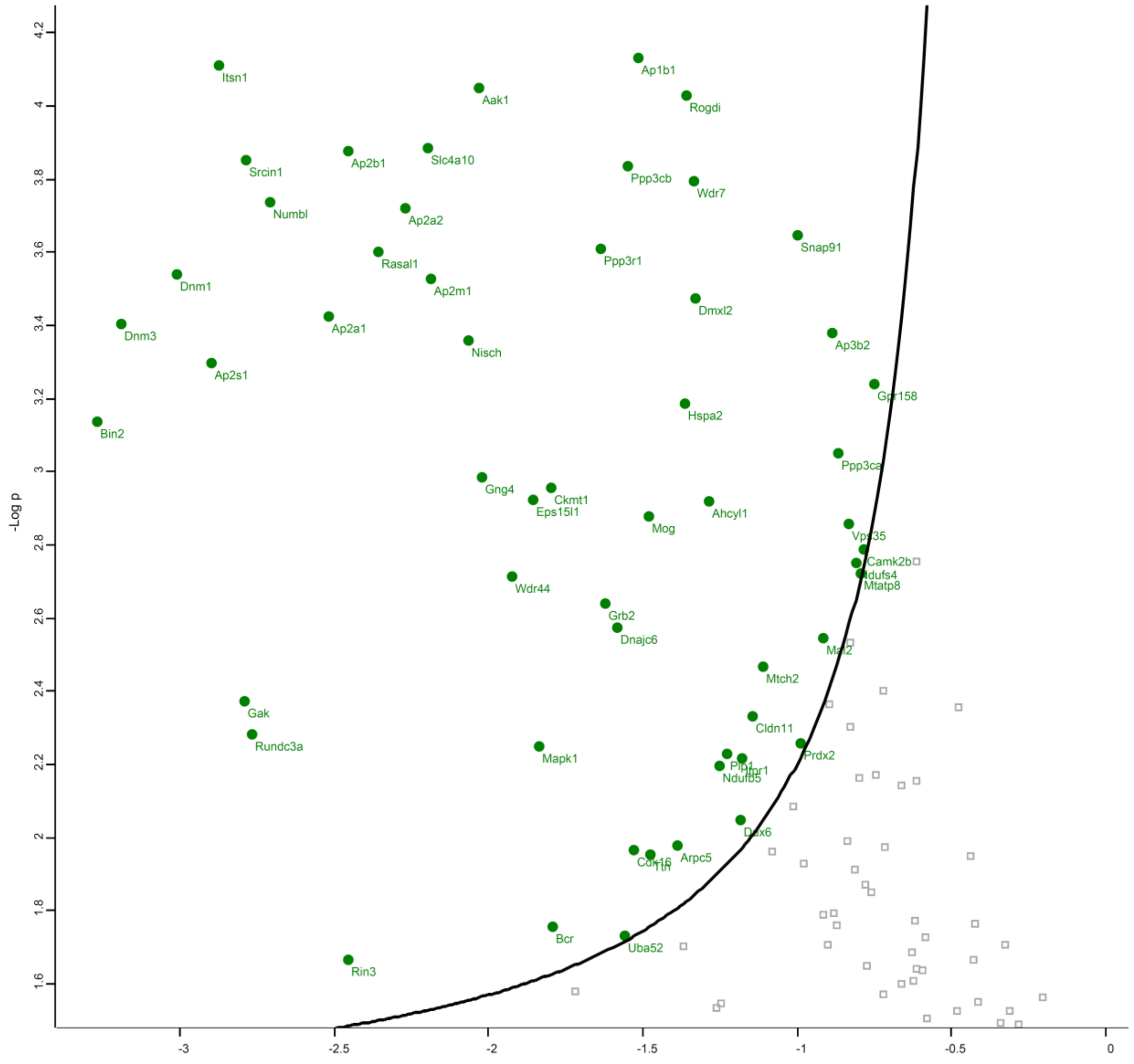


<u>Supplementary Fig. 13 </u>**. Proteins interacting with mouse BIN1 neuronal isoform**

Supplementary Fig. 13 . **Proteins interacting with mouse BIN1 neuronal isoform**

Volcano plot representation of 63 proteins statistically significant as a function of control immuno precipitations. Five transgenic brain of Tagged Knockin 3-old-month mice and five of WT C57B6 mice were used as described in Material & Methods.

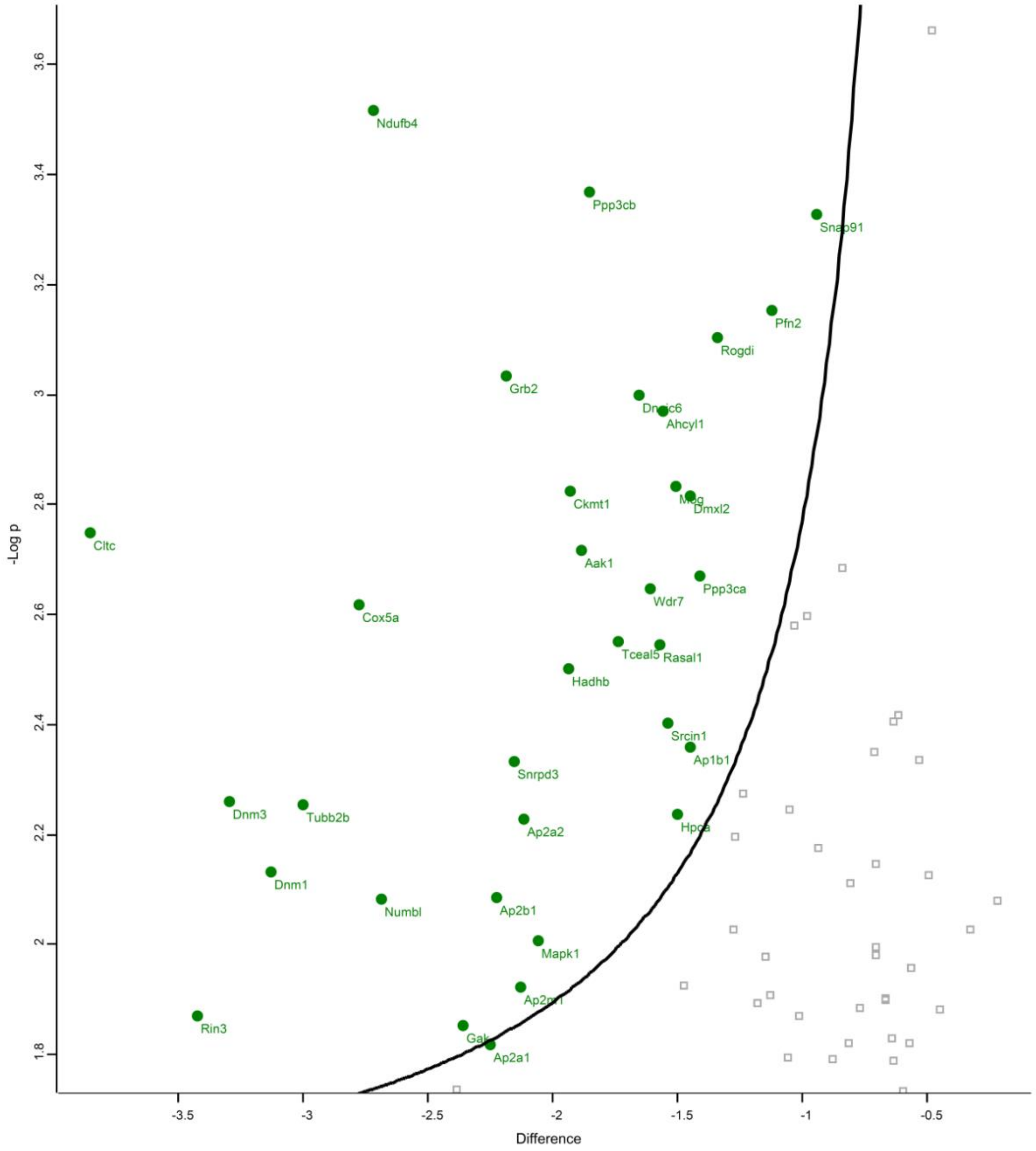


<u>Supplementary Fig. 14</u> . **Proteins interacting with all mouse BIN1 isoforms**

Supplementary Fig. 14. **Proteins interacting with all mouse BIN1 isoforms**

Volcano plot representation of 44 proteins statistically significant as a function of control immuno precipitations. Four transgenic brain of Tagged Knockin 3-old-month mice and five of WT C57B6 mice were used as described in Material & Methods.

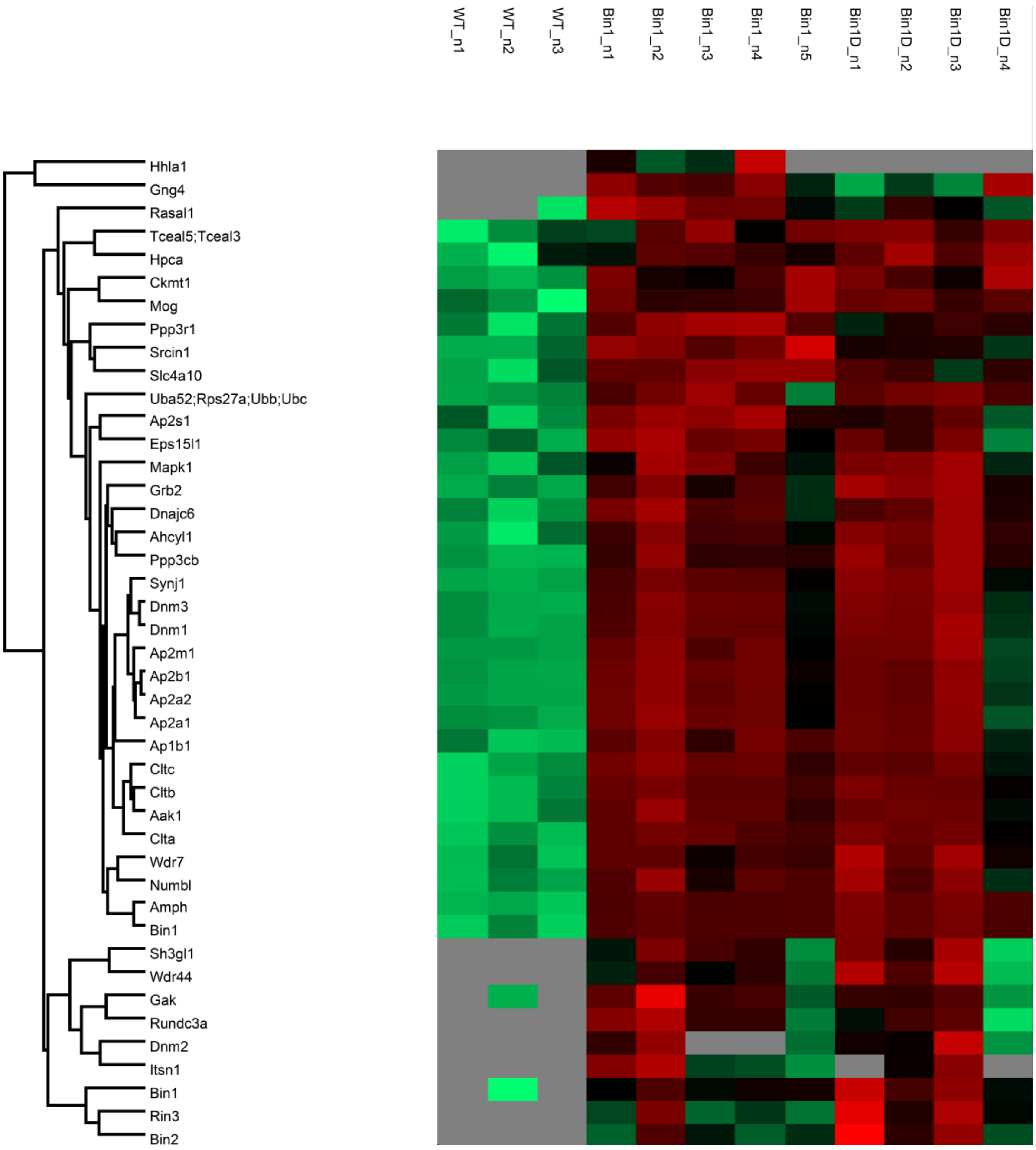


<u>Supplementary Fig. 15.</u> **Differential heat map of proteins identified in BIN1 immunoprecipitation complexes for tagged Bin1 knockin mice.**

Supplementary Fig. 15. **Differential heat map of proteins identified in BIN1 immunoprecipitation complexes for tagged Bin1 knockin mice.**

Differential heat map of proteins identified either in knockin Bin1 (Tag for neuronal isoform) or Bin 1D (C terminal Tag for all isoforms)**.** Note that for Bin1 (Tag for neuronal isoform) mice, RASAL1 was identified. RASAL1 is known to interact with bona fide post synaptic density proteins (DLG4, SYNGAP1, AGAP2) as shown in Li J, Zhang W, Yang H, Howrigan DP, Wilkinson B, Souaiaia T, Evgrafov OV, Genovese G, Clementel VA, Tudor JC, Abel T, Knowles JA, Neale BM, Wang K, Sun F, Coba MP. Spatiotemporal profile of postsynaptic interactomes integrates components of complex brain disorders. Nat. Neurosci. Aug. 01, 201

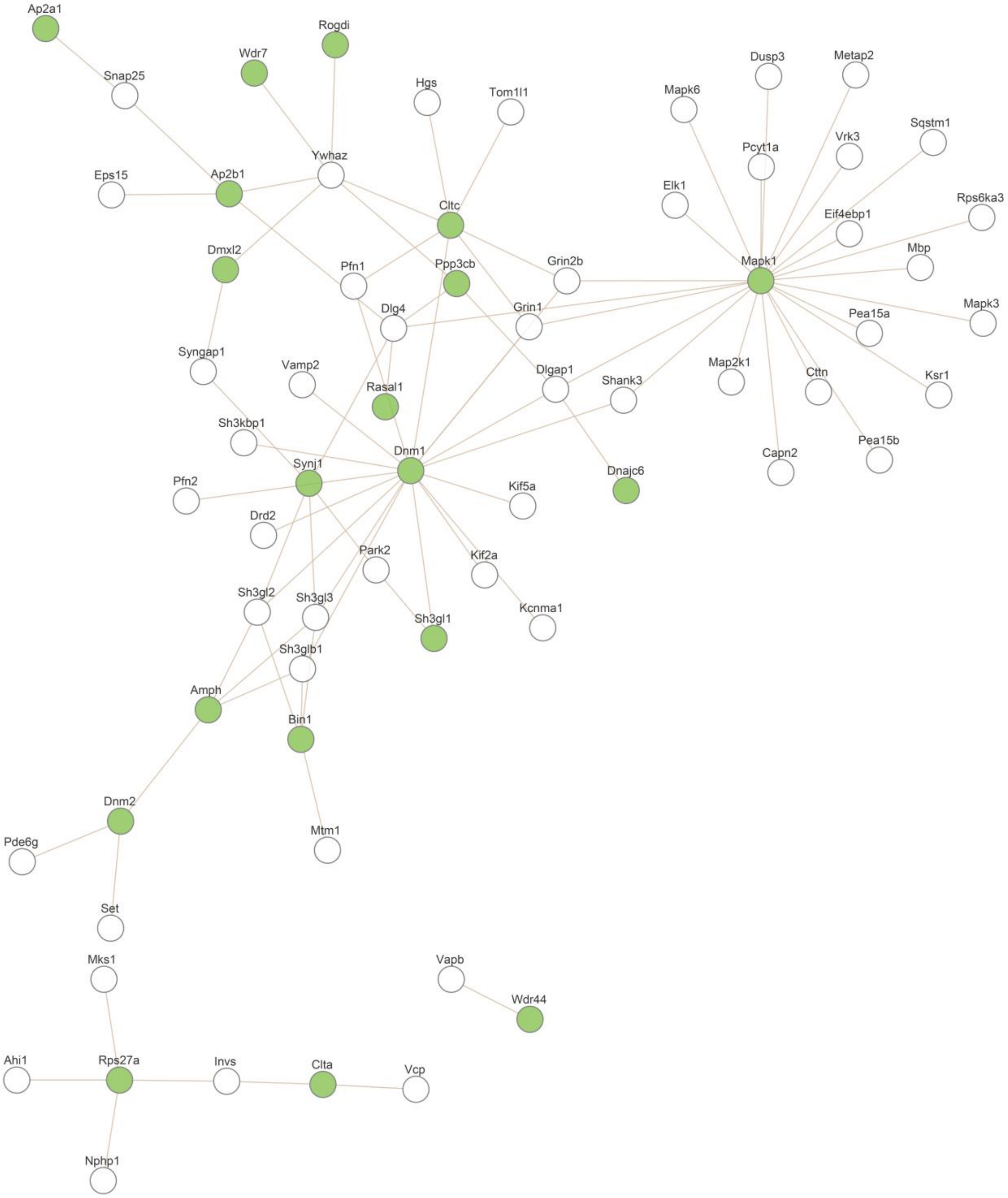


Supplementary Fig. 16. **Network analysis from Knockin mice: Chemical synaptic transmission network.**

Supplementary Fig. 16. **Network analysis from Knockin mice: Chemical synaptic transmission network.**

Module: PPI_BIOGRID_Module_49

ID:PPI_BIOGRID_M49

C=1008; O=19; E=6.79; R=2.8; PValue=5.38e-06; FDR=6.9e-04

Description:

This module was generated from m musculus BIOGRID PPI network (https://thebiogrid.org/download.php), which contains 1024 genes and 2060 edges.

Related Function:

Biological Process: chemical synaptic transmission (p-value < 2.220446e-16)

Cellular Component: endoplasmic reticulum (p-value < 2.220446e-16)

Molecular Function: cytoskeletal protein binding (p-value < 2.220446e-16)

We used Webgestalt 2017 suite analysis (Wang, J., Vasaikar, S., Shi, Z., Greer, M., & Zhang, B. WebGestalt 2017: a more comprehensive, powerful, flexible and interactive gene set enrichment analysis toolkit. Nucleic Acids Res. 2017 Jul 3;45(W1):W130-W137.

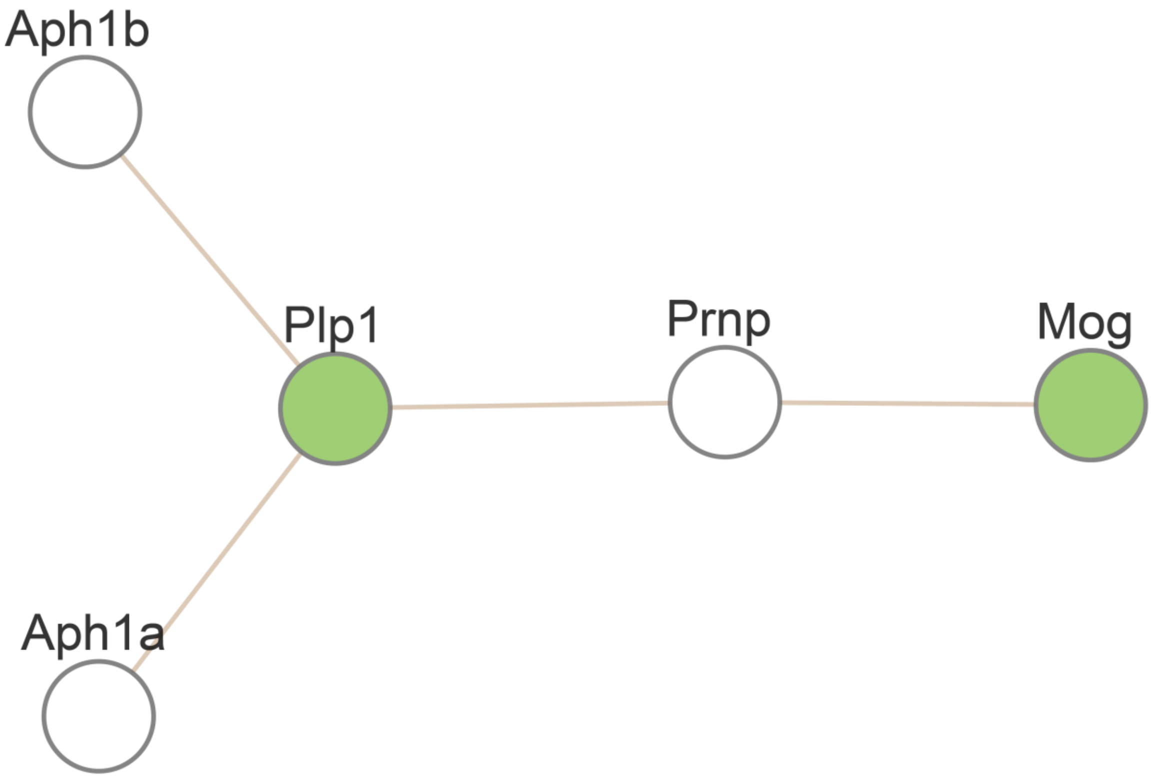


Supplementary Fig.17. **Network analysis from Knockin mice: Myelin sheath network.**

Supplementary Fig.17. **Network analysis from Knockin mice: Myelin sheath network.**

ID:PPI_BIOGRID_M52

C=33; O=2; E=0.22; R=9; PValue=2.05e-02; FDR=7.87e-01

Module: PPI_BIOGRID_Module_52

Description:

This module was generated from m musculus BIOGRID PPI network (https://thebiogrid.org/download.php), which contains 33 genes and 44 edges.

Related Function:

Biological Process: cellular metal ion homeostasis (p=7.04e-06)

Cellular Component: myelin sheath (p=4.56e-07)

Molecular Function: ATPase activity, coupled to transmembrane movement of ions, phosphorylative mechanism (p=9.23e-05)

We used Webgestalt 2017 suite analysis (Wang, J., Vasaikar, S., Shi, Z., Greer, M., & Zhang, B. WebGestalt 2017: a more comprehensive, powerful, flexible and interactive gene set enrichment analysis toolkit. Nucleic Acids Res. 2017 Jul 3;45(W1):W130-W137).

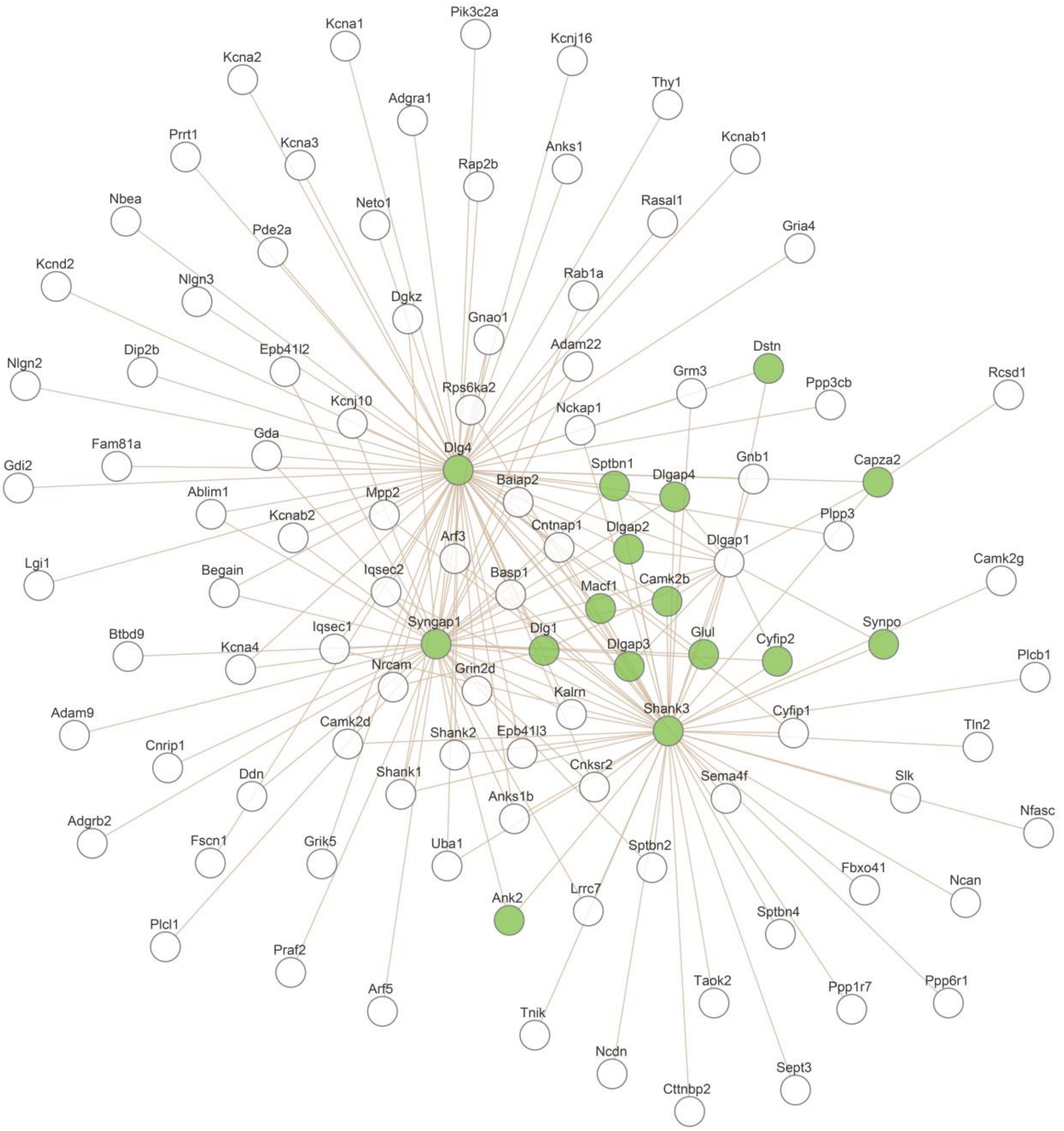


Supplementary Fig.18. **Network analysis from IP of C57B6 mouse synaptosomes: Chemical synaptic transmission network.**

Supplementary Fig.18. **Network analysis from IP of C57B6 mouse synaptosomes: Chemical synaptic transmission network.**

ID:PPI_BIOGRID_M49

C=1008; O=60; E=20.56; R=2.92; PValue=0e+00; FDR=0e+00

Module: PPI_BIOGRID_Module_49

Description:

This module was generated from m musculus BIOGRID PPI network (https://thebiogrid.org/download.php), which contains 1024 genes and 2060 edges.

Related Function:

Biological Process: chemical synaptic transmission (p-value < 2.220446e-16)

Cellular Component: endoplasmic reticulum (p-value < 2.220446e-16)

Molecular Function: cytoskeletal protein binding (p-value < 2.220446e-16)

We used Webgestalt 2017 suite analysis (Wang, J., Vasaikar, S., Shi, Z., Greer, M., & Zhang, B. WebGestalt 2017: a more comprehensive, powerful, flexible and interactive gene set enrichment analysis toolkit. Nucleic Acids Res. 2017 Jul 3;45(W1):W130-W137).

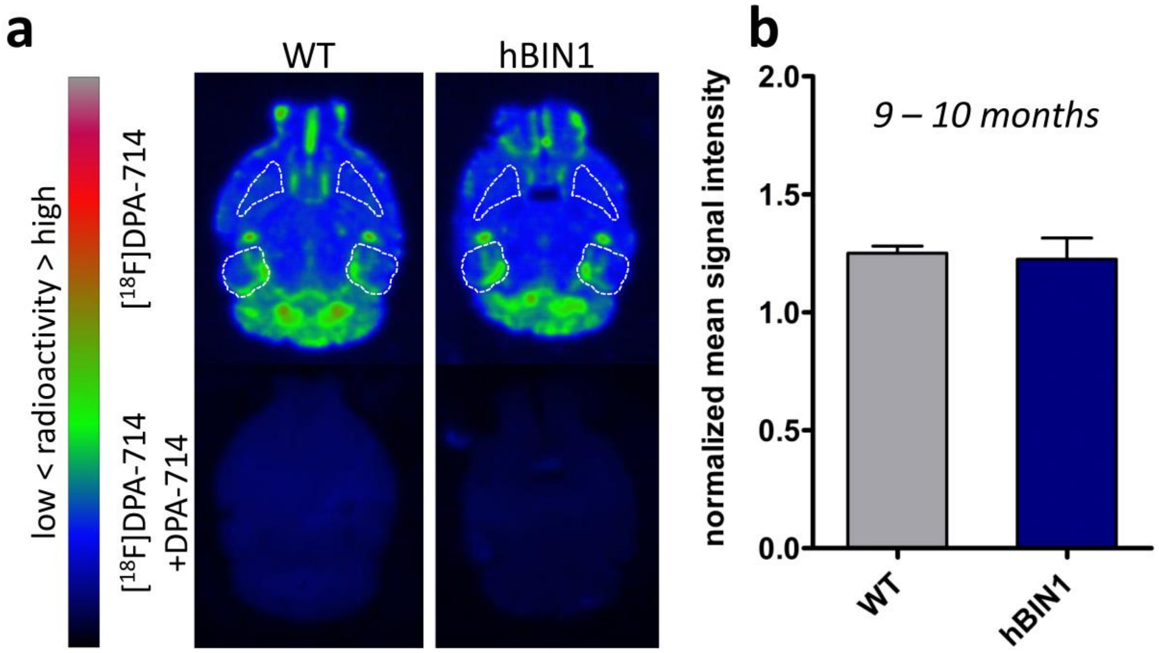


Supplementary Fig.19. **[$^{18}$F]DPA-714 quantification in hBIN1 and WT mice at month 15.**

Supplementary Fig.19. **[$^{18}$F]DPA-714 quantification in hBIN1 and WT mice at month 15.** Autoradiograms (**a**) and quantification (**b**) of axial brain sections from WT (n=3) and hBIN1 transgenic mice (n=3). Sections from WT and hBIN1 transgenic mice show some [$^{18}$F]DPA-714 binding in the dentate gyrus / medial entorhinal cortex region and only low binding in the striatum (**a, upper part**). Displacement with unlabeled DPA-714 demonstrates specific binding of [$^{18}$F]DPA-714 (**a, lower part**). The mean signal intensity for the dentate gyrus / medial entorhinal cortex region normalized to the striatum does not show significant differences for [$^{18}$F]DPA-714 binding between WT and hBIN1 transgenic mice (**b**), which may be due to changes between WT and transgenic animals that are not important enough to be detected considering the sensitivity of autoradiography using [$^{18}$F]DPA-714 and/or also due to the limitations of [$^{18}$F]DPA-714 quantification using Phosphor-Imager screens.
